# Revealing neural correlates of behavior without behavioral measurements

**DOI:** 10.1101/540195

**Authors:** Alon Rubin, Liron Sheintuch, Noa Brande-Eilat, Or Pinchasof, Yoav Rechavi, Nitzan Geva, Yaniv Ziv

## Abstract

Measuring neuronal tuning curves has been instrumental for many discoveries in neuroscience but requires a-priori assumptions regarding the identity of the encoded variables. We applied unsupervised learning to large-scale neuronal recordings in behaving mice from circuits involved in spatial cognition, and uncovered a highly-organized internal structure of ensemble activity patterns. This emergent structure allowed defining for each neuron an ‘internal tuning-curve’ that characterizes its activity relative to the network activity, rather than relative to any pre-defined external variable – revealing place-tuning in the hippocampus and head-direction tuning in the thalamus and postsubiculum, without relying on measurements of place or head-direction. Similar investigation in prefrontal cortex revealed schematic representations of distances and actions, and exposed a previously unknown variable, the ‘trajectory-phase’. The structure of ensemble activity patterns was conserved across mice, allowing using one animal’s data to decode another animal’s behavior. Thus, the internal structure of neuronal activity itself enables reconstructing internal representations and discovering new behavioral variables hidden within a neural code.

---

Most neurons in the brain do not receive direct inputs from the external world; rather, their activity is governed by their interactions with other neurons within and across brain circuits. Despite this fact, studies in neuroscience typically focus on neuronal responsiveness to an examined external variable (i.e., a *neuronal tuning curve)*. Rooted in the emergence in the 1950’s of electrophysiological techniques for recording from single neurons in vivo, this ‘neural correlate’ approach opened the door to studying how specific brain circuits form internal representations, and has led to seminal breakthroughs. Examples of such breakthroughs include the discoveries of orientation tuning in the visual cortex^1^, hippocampal place cells^2^, and entorhinal grid cells^3,4^. However, while such analyses remain invaluable for many neuroscientific studies, they are limited to a-priori defined external variables, overlooking other variables that were not measured, considered relevant, or observable^5–7^. With traditional electrophysiological techniques that permitted recordings from only small numbers of neurons, no feasible alternatives to this approach existed. Recent advances in multi-electrode and optical imaging technologies enable simultaneous readout of activity from large neuronal populations, permitting a qualitatively different approach to study the neural code—via the attributes of neuronal activity itself.

We hypothesized that the relationships between neuronal population activity patterns would give rise to a structure within the neuronal activity space that differs across brain circuits according to their distinct computational roles. Thus, studying the internal structure of neuronal activity itself could reveal key properties of the neural code that are specific to a given brain circuit, without relying on measurements of the behavior or the stimulus. We further hypothesized that characterizing the activity of single neurons with respect to this structure (rather than relative to any pre-selected external variable), could yield *internal tuning curves*, which capture the properties of classically-computed tuning curves. Realization of such an approach could enable the investigation of neural coding in brain circuits for which little is known about the identity of the variables they encode, and circumvent biases associated with the naïve application of the neural correlate approach.

## Results

### Internal representation of space in the hippocampus

First, we sought to study a brain circuit that is known to encode a canonical variable, and test whether its coding properties could be extracted from the internal structure of neuronal activity. Thus, we focused on the dorsal CA1 of the hippocampus, a circuit in which many neurons are tuned to spatial position^2^. We used miniaturized head-mounted microscopes^8^ to image Ca^2+^ dynamics in GCaMP6 expressing hippocampal CA1 neurons in freely behaving mice. During imaging, mice ran back and forth along a linear track to collect water rewards^9^ (**Fig. 1a**). After detecting cells and Ca^2+^ events in the imaging data^9,10^, we constructed neuronal ensemble activity vectors of instantaneous neuronal activity sampled at fixed time bins. To explore the relationships between the ensemble activity patterns we applied a non-linear dimensionality reduction algorithm (Laplacian Eigenmaps^11^; LEM; **Supplementary Fig. 1**) to the activity vectors (**Fig. 1b** and **Supplementary Movie 1**). This analysis revealed a structure in the reduced dimensional neuronal activity space, with a high density of data points within a small number of clusters (clusters A, B, C, D, and E in **Fig. 1b**), suggesting that throughout the imaging session, the network leaped between discrete states. Temporally segmenting the data points into time intervals during which the network remained within a given state, revealed a recurring cyclic pattern of transitions between network states (**Fig. 1c**). Within a cycle, each state appeared only once, except for state C, which appeared twice, in two different phases. Further analysis revealed that the set of segments of state C consisted of two separate state subtypes (C_1_ and C_2_; **Fig. 1d**), each corresponding to one of the two different phases (red and blue segments in **Fig. 1c**). These analyses allowed us to portray the intricate structure of network states and their pattern of transitions (**Fig. 1e-f**).

**Fig. 1:**
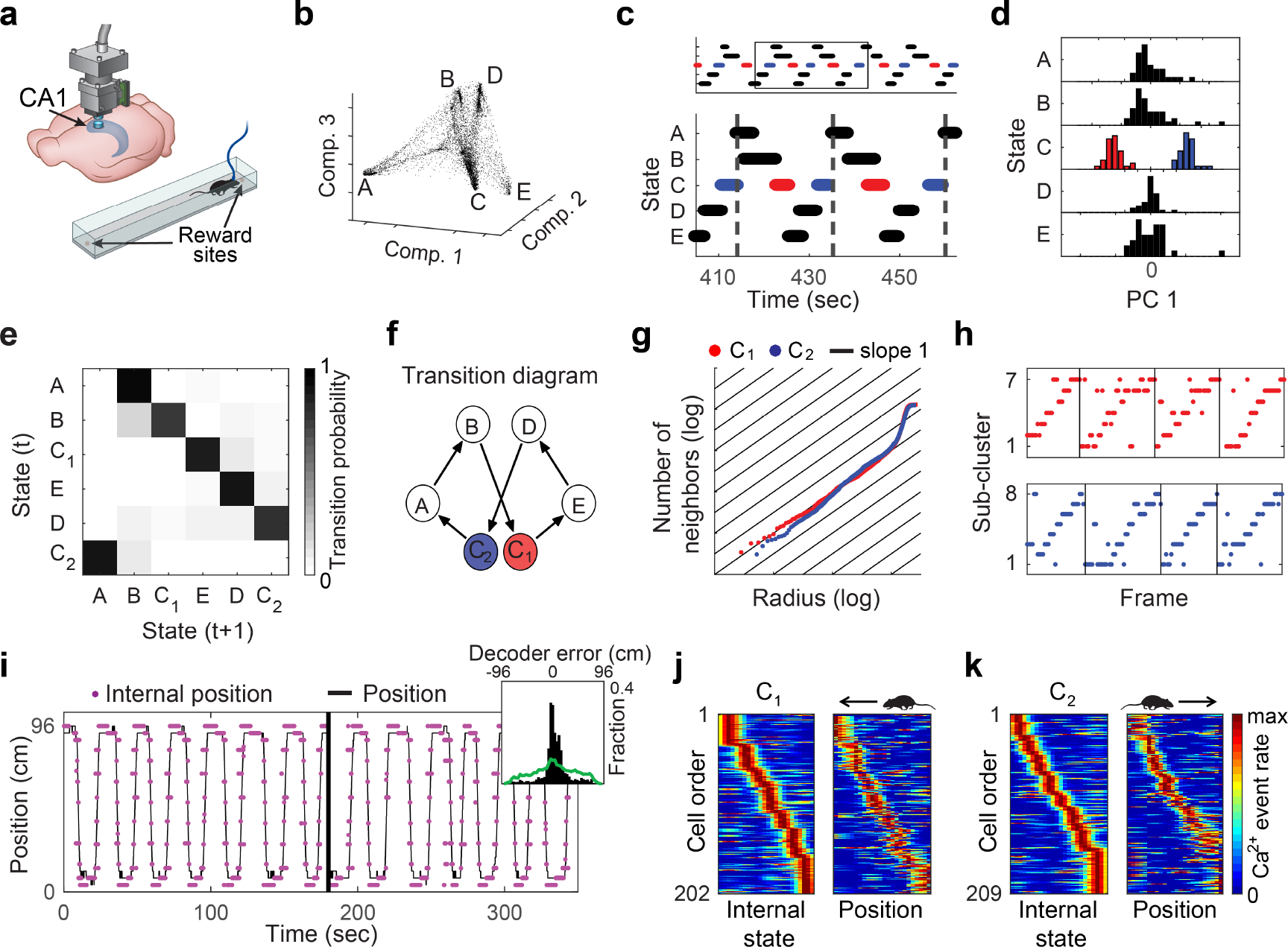
Hippocampal internal representation of position obtained without relying on behavioral data. (**a**) Ca^2+^ imaging using miniaturized head-mounted microscopes in the hippocampal CA1 of freely behaving mice. Mice ran back and forth to collect rewards at the two ends of a linear track. (**b**) The distribution of data points in the reduced dimensional space of neuronal activity forms a small number of dense clusters. (**c**) The temporal structure of segmented data reveals a cyclic pattern of transitions between network states. Each state type appeared once per cycle except for state C which appeared in two different phases. (**d**) Principal component analysis applied separately to the segment-level activity patterns of each state indicated that only state C consists of two completely separate clusters (red and blue, subtypes C_1_ and C_2_, respectively). Histogram of the projections on the first principle component is presented for each state. Note that the two subtypes consistently appear in two different phases in the cyclic temporal pattern, as shown in **c**. (**e**) The segment-level transition matrix (i.e., the probability of a segment to be in cluster i, given that the preceding segment is in cluster j) shows a stereotypical pattern of transitions between the different network states. (**f**) Illustration of the structure of network states and the stereotypic pattern of transitions between them. (**g**) Estimation of internal dimension calculated separately for data taken from segment subtypes C_1_ (red) and C_2_ (blue): Cumulative number of neighboring data points as a function of the radius in the reduced dimensionality space, plotted on a log-log scale. The slope of the data is close to one (black lines), indicating a dimension of one. (**h**) Sub-clusters within segments of subtype C_1_ (top) and C_2_ (bottom) exhibited a stereotypic temporal structure. Different trajectories within segments of subtype C_1_ and C_2_ are separated by black vertical lines indicate (**i**) The reconstructed internal position (magenta) and the actual position (black) of the mouse while freely exploring the linear track. Inset, distribution of the error in the reconstruction of position (black) versus shuffled data (green). (**j**-**k**) For each neuron, we calculated its internal tuning curve, relative to the internal state of the network (left), and the external tuning curve, relative to the position of the animal, namely place fields (right). Neuronal activity within segment subtype C_1_ is presented next to the activity of the same cells during leftward running (**j**), and activity within segment subtype C_2_ is presented next to the activity of the same cells during rightward running (**k**).

By sorting the behavioral data according to the specific network states, we found that each state corresponded to a different behavior and location along the linear track (**Supplementary Movie 2**). Specifically, states A and E, which were symmetrically located within the internal structure (**Fig. 1b**), corresponded to drinking at the left and right sides of the track, respectively. Similarly, the symmetric states B and D corresponded to turning at the two sides of the track. States C_1_ and C_2_ corresponded to epochs of running in different directions along the linear track, consistent with the characteristics expected for spatial coding in one-dimensional environments^12,13^. Thus, the internal structure of neuronal activity exposed discrete sets of network states that corresponded to different combinations of locations and behaviors, without needing to a-priori hypothesize that these specific behavioral states are encoded by hippocampal neurons.

We next attempted to use the structure of the neuronal activity to characterize the internal representation of space at a finer resolution, and focused our analysis on states C_1_ and C_2_. We applied the dimensionality reduction procedure separately for the data within C_1_ and C_2_. Then, we estimated the internal dimension of the data14 (**Supplementary Fig. 2**) of each state subtype and found that both C_1_ and C_2_ were one-dimensional (**Fig. 1g**). By sub-clustering the data points within C_1_ and C_2_, we found an ordered pattern of transitions between the sub-clusters throughout each segment (**Fig. 1h**), which monotonically covered a one-dimensional continuum of network states. Consistent with these observations, the trajectory of the network within the internal structure of neuronal activity reflected the trajectory of the mouse along the track, permitting accurate reconstruction of position (**Fig. 1i**; permutation test, p<0.001 for each mouse, N=4). Note that because the reconstructed trajectory is not calibrated to the internal symmetry of the encoded variable nor to its identity, we set the reflection degree of freedom of the internal representation to match the position of the mouse. We then calculated the internal tuning curves of individual neurons: i.e., the activity of single neurons with respect to the different network states. The internal tuning curves had similar properties to those of their corresponding external tuning curves (i.e., place fields) (**Fig. 1j-k** and **Supplementary Fig. 3a**; permutation test, p<0.001 for each mouse, N=4). These results demonstrate that the relationships among neuronal activity patterns themselves are sufficient to reconstruct the hippocampal representation of space.

### Internal representations in the medial prefrontal cortex

Thus far, our analysis focused on data recorded from the hippocampus, a brain region in which the encoding of a known variable (place) is dominant. We next examined whether a similar analysis can reveal internal representations using recordings from brain circuits that have not been associated with a canonical encoded variable. To this end, we focused on the anterior cingulate cortex (ACC), a sub-region within the medial prefrontal cortex (mPFC) that is involved in multiple high-order cognitive processes^15–18^. We conducted Ca^2+^ imaging in the ACC of freely behaving mice performing the same linear track exploration task described above for hippocampal imaging (**Fig. 2a**). Dimensionality reduction of the neuronal activity revealed a high density of data points within a small number of clusters (**Fig. 2b**), similar to our observations in the hippocampus. Next, we clustered the data points into different network states and calculated their transition matrix (**Fig. 2c**), which exhibited a recurring cyclic pattern (**Fig. 2c-d**). In contrast to our observations from the hippocampal data, in the ACC the representation of behaviors on both sides of the track converged onto the same network states (**Fig. 2d**). Consistent with this observation, the distribution of the animal's position along the linear track was symmetrical for each of the identified network states (**Fig. 2e**). Furthermore, by sorting the behavioral data according to the network states, we found that each state corresponded to a distinct behavior, irrespective of the position of the mouse. We termed these behaviors ‘Rearing’, ‘Turning’, ‘Start run’, ‘End run’, and ‘Drinking’ (**Supplementary Movie 3**). We confirmed this clustering-based classification using manual labeling of the mouse behavior (**Fig. 2f**).

**Fig. 2:**
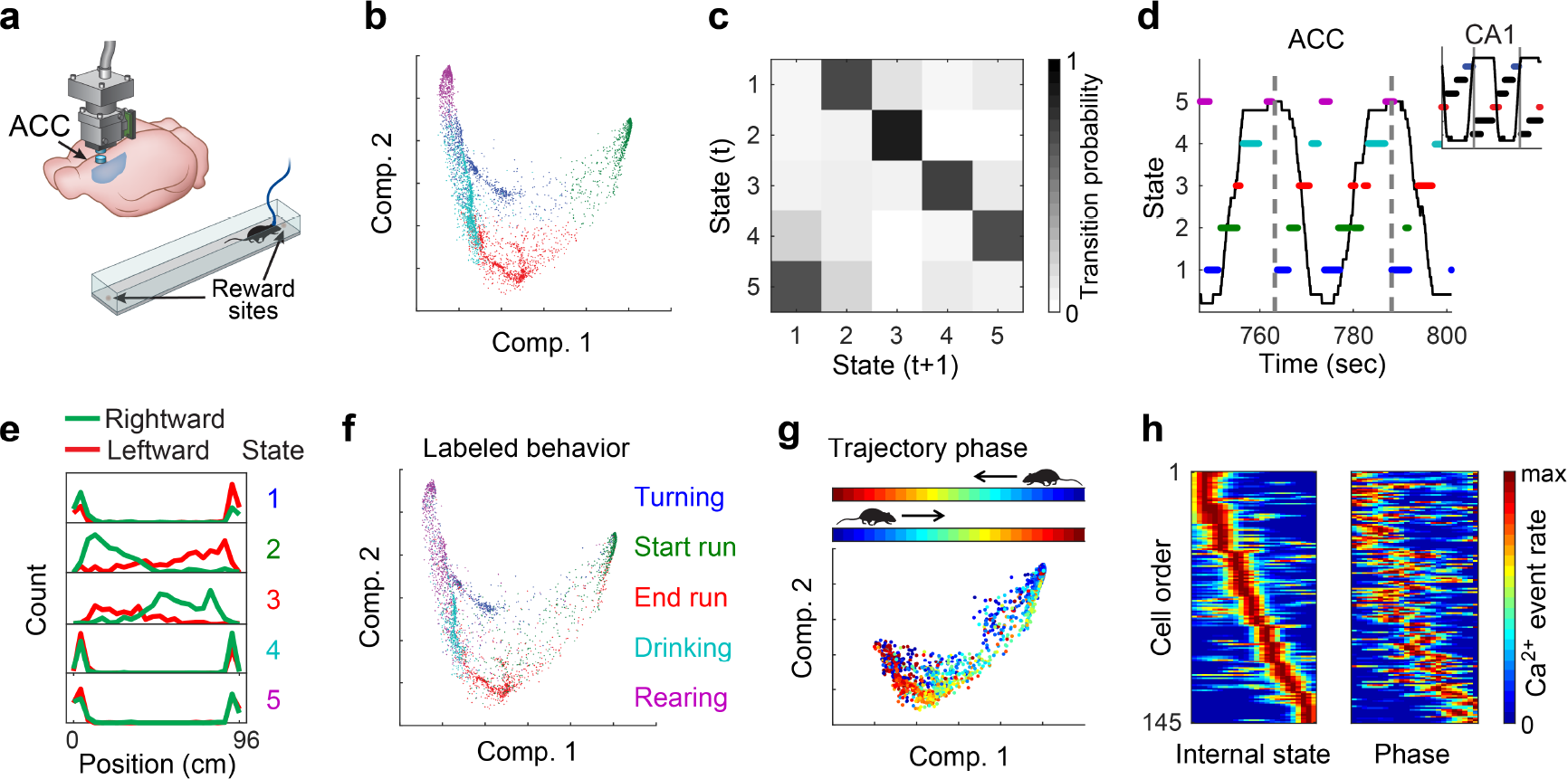
ACC activity reveals a schematic representation of behavior along the linear track. (**a**) Ca^2+^ imaging using miniaturized head-mounted microscopes in the ACC of freely behaving mice. Mice ran back and forth to collect rewards at the two ends of a linear track. (**b**) The distribution of data points in the reduced dimensional space of neuronal activity. The data are clustered and colored to indicate each of the different clusters. (**c**) The segment-level transition matrix shows a stereotypical pattern of transitions between the different network states. (**d**) A stereotypic temporal structure of network states (marked by different colors) is observed during exploration of the linear track (animal position is shown in black). A full running cycle (bounded between vertical gray lines) corresponds to two cycles of transitions between network states. Inset, hippocampal data, in which a full running cycle (bounded between vertical gray lines) corresponds to a single cycle of transitions between network states. (**e**) The distribution of mouse position along the linear track is symmetrical for each of the network states. (**f**) The same distribution presented in **a**, with data points colored according to behavioral labeling. (**g**) Distribution of data points in the reduced dimensional space of neuronal activity for the two clusters corresponding to running. Data points are colored according to trajectory phase, i.e., the distance of the mouse from the start of the track (opposite end for each running direction). (**h**) For each neuron, we calculated its internal tuning curve, relative to the internal state of the network (left), and the external tuning curve, relative to the trajectory phase of the animal (right).

To study the structure of the neuronal activity in the ACC at a finer resolution, we focused on the network states that were linked to running (‘Start run’ and ‘End run’). We found that neuronal activity represented the position of the mouse relative to the start and end points of each track traversal, regardless of the running direction (i.e. the trajectory phase; **Fig. 2g**). Consistent with this observation, constructing internal tuning curves revealed individual neurons that were tuned to a specific phase, namely, trajectory-phase cells (**Fig. 2h** and **Supplementary Fig. 4a-b**). The internal tuning captured the external tuning of the same neurons to the trajectory phase (**Fig. 2h** and **Supplementary Fig. 3b**; permutation test, p<0.001 for each mouse, N=3). The observed encoding of trajectory phase could not be accounted for by velocity or acceleration (**Supplementary Fig. 5**). Thus, even in a brain circuit in which less is known about the identity of the encoded variables, our analysis exposed key properties of the internal representation, and the encoding of a previously unknown variable.

### Different internal structures of neuronal activity in the ACC and the hippocampus reflect different types of representations of locations and actions

Having found different structures of neuronal activity in the ACC and hippocampus under the same behavioral task, we sought to further characterize the differences between their internal representations of locations and actions. Neuronal activity in the ACC, but not in the hippocampus, was similar during epochs of the same behavioral state at opposite sides of the track (**Fig. 3a**, and **Supplementary Fig. 6**; one sample t-test, p>0.05 and p<0.05 for each behavioral state for CA1 and ACC, respectively). This distinction between the neuronal coding in the hippocampus and ACC was also evident when examining the tuning of individual cells (**Fig. 3b** and **Supplementary Fig. 4**). To further substantiate the differences between the encoding of trajectory phase and position, we devised a “phase decoder”, and found that in the ACC we could accurately infer the trajectory phase in a given direction, when the decoder was trained on the relationship between neuronal activity and the trajectory phase in the opposite direction (**Fig. 3c**; permutation test, p<0.001 for each mouse, N=3). Notably, we could not predict the trajectory phase when applying the same analysis to hippocampal data (**Fig. 3c**; permutation test, p>0.05 for each mouse, N=4). Consistent with these findings, population activity in the ACC was correlated between symmetrical spatial locations for opposite running directions along the linear track (**Fig. 3d-e**). In contrast, in the hippocampus, neuronal representations differed considerably between the two sides of the track, and were more sharply tuned to position than in the ACC (**Fig. 3f**). Overall, we found that ACC neurons are spatially tuned, but the nature of their spatial representation is markedly different from the classical spatial tuning properties of hippocampal neurons.

**Fig. 3:**
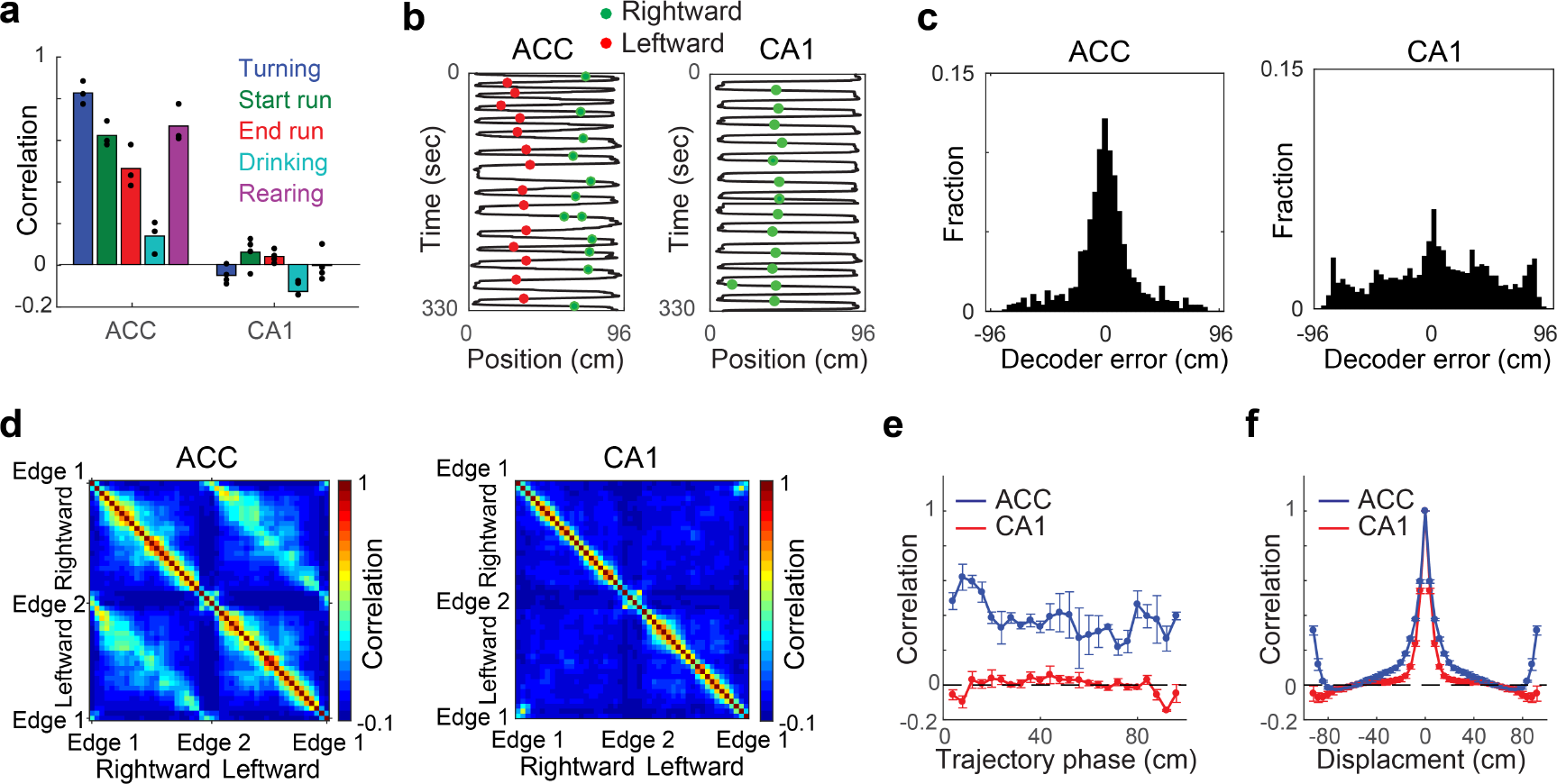
The different internal structures in the ACC and the hippocampus reflect different representations of locations and actions. (**a**) Pearson correlation between ensemble activity patterns from the two sides of the linear track, for each behavioral state, for data recorded in the ACC (left), and the hippocampus (right). Ensemble activity patterns consist of concatenated epochs from a given behavioral state on the same side of the track. Data show means for N=3 mice in the ACC (left), and N=4 mice in the hippocampus (right). (**b**) An example trajectory-phase cell recorded in the ACC (left), and an example place cell recorded in the hippocampus (right). Black lines show the positions of the animals, and the green and red dots show activity of the neurons during rightward and leftward running, respectively. (**c**) Distribution of decoding error of trajectory phase for data recorded in the ACC (left), and the hippocampus (right). The decoder was trained on data from running in one direction and tested on data from running in the other direction. (**d**) Pearson correlation between ensemble activity patterns from different spatial locations on the linear track, for data recorded in the ACC (left; averaged over N=3 mice), and the hippocampus (right; averaged over N=4 mice). Ensemble activity patterns consist of concatenated epochs from a given location, separated according to the two running directions. (**e**) Pearson correlation between ensemble activity patterns from the two sides of the linear track, given the same trajectory phase, for data recorded in the ACC (blue), and the hippocampus (red). Ensemble activity patterns are defined as the mean event rate for each neuron given a spatial bin and running direction. (**f**) Pearson correlation between ensemble activity patterns for different spatial displacements on the linear track, for data recorded in the ACC (blue), and the hippocampus (red). Correlations are averaged over mice and over the two running directions. Data in **e-f** show means ± SEM, for N=3 mice in the ACC (blue), and N=4 mice in the Hippocampus (red).

### Internal representation of head direction during wake and REM sleep periods

Since our approach does not rely on behavioral measurements, we next asked if we can use it to expose internal representations even when there is no correspondence between neuronal activity and the external stimulus or behavior. It has been shown that pairwise correlations between head direction neurons in the anterior dorsal nucleus of the thalamus (ADn) and the postsubiculum (PoS) are preserved during sleep^19^. Therefore, we analyzed published electrophysiological recordings from the ADn and the PoS^19,20^ in mice foraging for food in open environments (**Fig. 4a**), and sought to compare the internal structure of neuronal activity during wake and sleep periods. We constructed neuronal ensemble activity vectors as described above for the Ca^2+^ imaging data, and applied dimensionality reduction to the activity vectors, separately for wake periods and periods of REM sleep. This analysis exposed a ring structure in the reduced dimensional space of neuronal activity^21,22^ for both wake and REM data (**Fig. 4b-c** and **Supplementary Fig. 1c-j**). By quantifying the internal dimension^14^ and topology^23–26^ of the data (**Supplementary Fig. 7a-d**), we found a one-dimensional structure (**Supplementary Fig. 8h**), with one component, one hole, and no spaces (**Supplementary Fig. 7e**,and **Supplementary Fig. 9i**), providing an additional indication that the encoded variable is indeed characterized by a ring topology both during wake and REM sleep periods. These properties are consistent with the encoding of head direction previously observed in the ADnand PoS^19,27,28^.

**Fig. 4:**
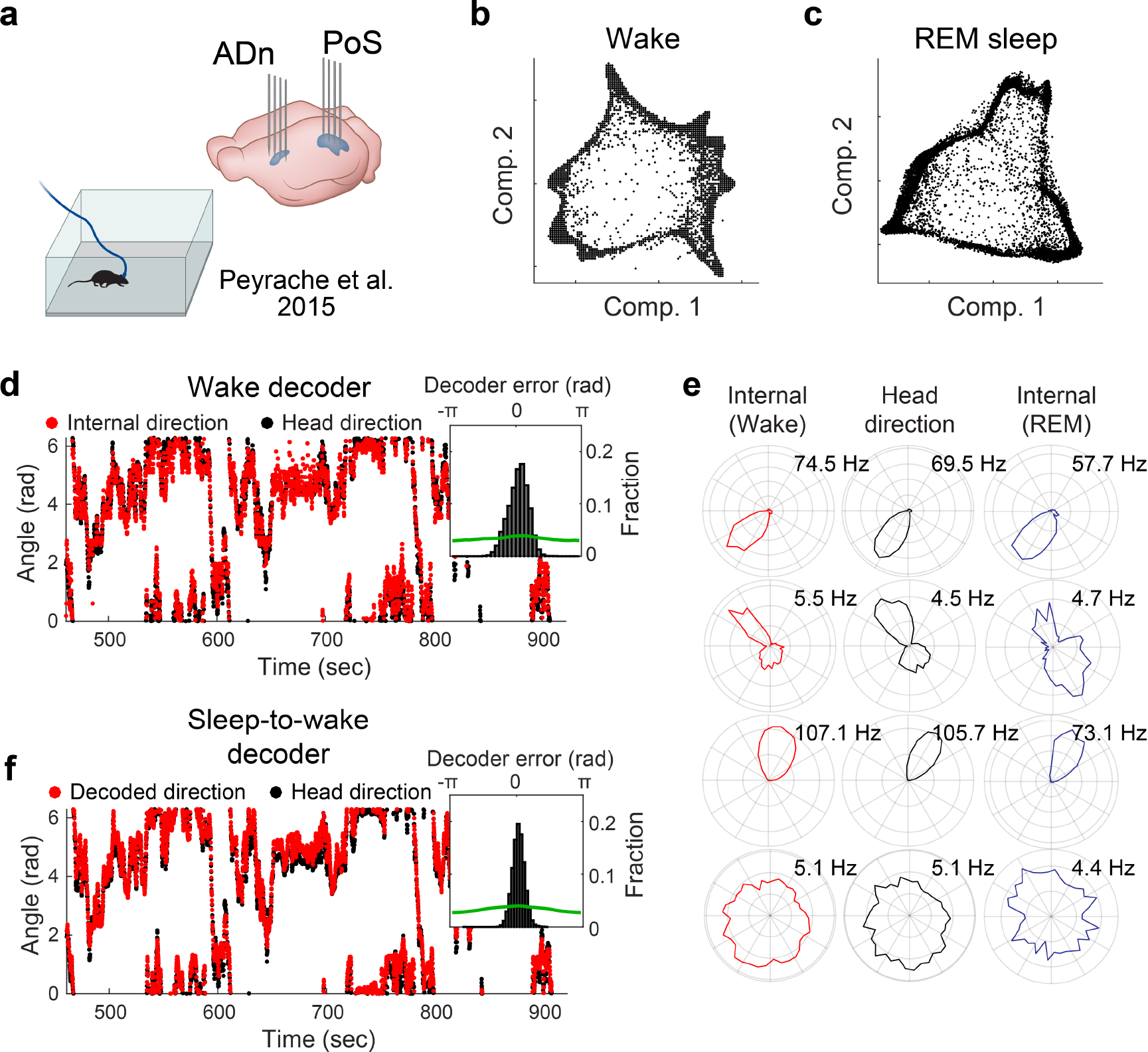
Internal representation of head direction during wake and REM sleep periods. (**a**) Data were obtained from dense electrophysiological recordings in the ADn and PoS, in mice foraging for food in open environments. (**b-c**) The distribution of data points in the reduced dimensional space of neuronal activity is characterized by a ring topology, during wake periods (**b**) and during periods of REM sleep (**c**). Each data point corresponds to the neuronal activity within a single time frame. (**d**) The reconstructed internal direction (red) and the actual head direction (black). Inset, distribution of the error in the reconstruction of head direction (black), versus shuffled data (green). (**e**) Internal tuning curve, relative to the state of the network (left, red), external head direction tuning curve, relative to mouse behavior (center, black), and the internal tuning curves during periods of REM sleep (right, blue) for four representative cells. (**f**) The decoded head direction (red) and the actual head direction (black). Inset, distribution of the decoding error (black) versus shuffled data (green). The decoder is based on internal tuning curves obtained exclusively from sleep.

Based on the temporal contiguity of the network states (**Supplementary Fig. 8a-b**), we reconstructed the trajectory of the network within the neuronal activity space and validated it against the head direction of the mouse. This analysis confirmed that the reconstructed trajectory reflected the actual head direction, up to the internal symmetries of a ring to reflection and rotation (**Fig. 4d**, **Supplementary Fig. 8c-d**, and **Supplementary Movie 4**). By accounting for the activity of each neuron relative to the network activity states of the entire recorded population, we calculated the internal tuning curves, and found that they were similar to the external tuning curves (**Fig. 4e**, **Supplementary Fig. 8e-g**). By applying a similar analysis to the data recorded during REM sleep, we found that the internal tuning curves obtained exclusively during REM matched the head direction tuning curves during wake periods (**Fig. 4e**, **Supplementary Fig. 9c-g**, and **Supplementary Movie 5**). Remarkably, based on the internal tuning curves that were obtained during REM sleep, we were able to accurately decode the head direction (up to the ring’s internal symmetry) while the mouse was awake and freely behaving (**Fig. 4f**). These observations suggest that in certain brain circuits the internal structure of neuronal activity is conserved irrespective of the animal’s behavior.

### Conservation of the internal structure of neuronal activity over days and across individuals

If the internal structure of neuronal activity reflects the computational processes undertaken by the network, then this structure should be an invariant property. To test this, we imaged neuronal activity in the hippocampus of mice that visited the same linear track on multiple days and found that the structure of neuronal activity remained similar over time (**Fig. 5a**), despite the substantial turnover of the active cells^9,29^ (41-64% overlap across days). The internal structure of neuronal activity in the hippocampus (**Fig. 5b**) and in the ADn and PoS (**Fig. 5c**) was also similar across different mice. To assess the similarity of the internal structures between mice and over time, we sought to identify analogous activity patterns across data sets (from different mice or different days), and test whether these patterns are also associated with similar behaviors. We devised an across-mice or across-days decoder (**Fig. 5d**), and found that we could accurately infer the position (**Fig. 5e-f** and **Supplementary Movie 6**) or head direction (**Fig. 5g**) of a mouse, based on the mapping between the behavior and activity patterns in another mouse or another day. Overall, these results demonstrate the conservation of internal structures of neuronal activity over time and across mice.

**Fig. 5:**
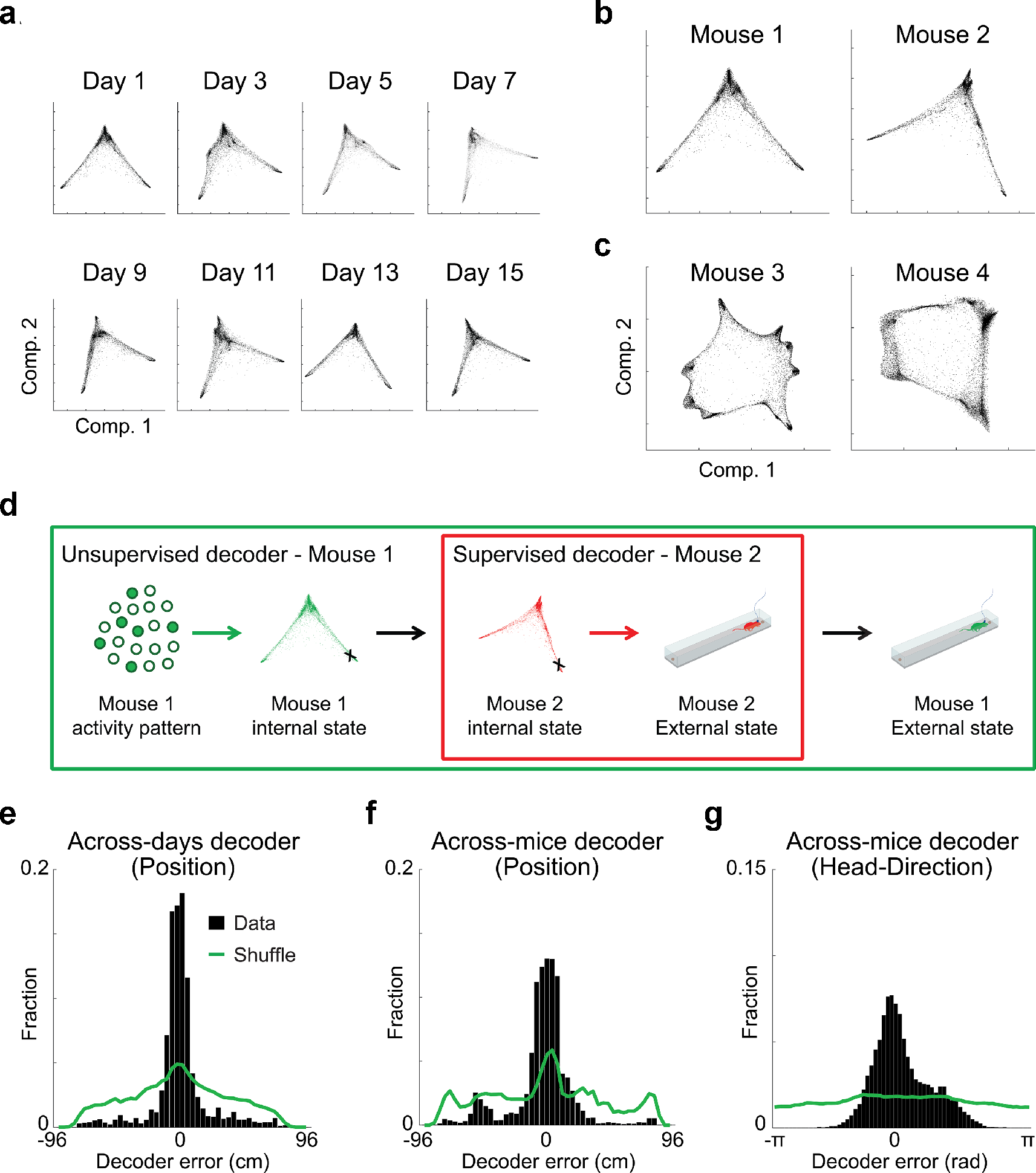
The internal structure of neuronal activity is maintained across days and across mice. (**a**) The distribution of data points in the reduced dimensional space of neuronal activity for hippocampal data from eight different days of the experiment. Note that the distribution of data points maintains a similar structure across days. (**b-c**) The distribution of points in the reduced dimensional space of neuronal activity for hippocampal data (**b**), and ADn/PoS data (**c**) from two different mice. (**d**) Workflow for the across-mice decoder. Neuronal activity in mouse 1 is associated with a specific network state within its internal structure of neuronal activity. Based on the similarity between internal structures, an analogous network state is found in mouse 2. This network state of mouse 2 is associated with a known external state. The decoded external state of mouse 1 is set as the same associated external state of mouse 2. (**e-g**) Distribution of decoding error for the across-days decoder for hippocampal data (**e**), the across-mice decoder for hippocampal data (**f**), and the across-mice decoder for ADn/PoS data (**g**).

## Discussion

Here, we introduced a new approach for studying the neural code based on the attributes of neuronal activity itself. By applying dimensionality reduction to large-scale neuronal data, we show that the internal structure of neuronal activity allows revealing key properties of the neural code in a given brain circuit. Previous studies have used dimensionality reduction methods to explore population-level coding of variables of interest in high dimensional neural data^5,30–36^. Unlike previous applications of dimensionality reduction in neuroscience, we demonstrate that internal representations and tuning curves can be reconstructed from the internal structure of neuronal activity, and that such reconstructions can be achieved even in brain circuits where the encoded variable is unknown. While in this work we used LEM for dimensionality reduction, it is likely that the approach we introduced here could accommodate other dimensionality reduction methods as well. Our analysis suggests, however, that non-linear dimensionality reduction methods are more suitable for studying the internal structure of neuronal activity, as they allow extracting a structure that accurately reflects internal representations in cases that linear methods fail (**Supplementary Fig. 1**).

Since our approach relies on minimal prior assumptions, and is applied irrespective of the specific identity of the encoded variables, it could circumvent biases and limitations that are inherent to the standard method of calculating neuronal tuning curves. Furthermore, calculating internal tuning curves could alleviate the need to set non-adaptive and arbitrary boundaries between different behavioral states when calculating (external) tuning curves. For example, place fields are typically calculated after applying position and velocity thresholds to define periods of locomotion^29,37^, and only neuronal activity during those periods is considered in the analysis. Our results demonstrate that neuronal activity itself can be used to identify behavioral states and the boundaries between them, enabling analysis that is derived explicitly from the neuronal activity data.

We used our approach to study internal representations in the hippocampus and the ACC while mice were performing the same behavioral task. Although both brain circuits exhibited a discrete set of network states that corresponded to different locations and behaviors, their internal structures of neuronal activity differed, reflecting different types of internal representations. The hippocampus represented a combination of locations and actions, and these representations were different between opposite sides of the track. In contrast, our analysis of recordings from the ACC exposed schema-like representations of distances and actions that are similar across the opposite sides of the track, including the encoding of a previously unknown variable – the trajectory phase. Previous work demonstrated that the mPFC is important for the assimilation of newly acquired information against prior knowledge, suggesting an underlying schematic organization of information^38^. Our findings are consistent with this notion, and specify how such schemas can be realized at the neural code level.

By studying internal representations of head direction in the ADn and PoS, we demonstrated that the internal structure of neuronal activity can reveal internal representations even when there is no correspondence between neuronal activity and the external stimulus or behavior (e.g., during sleep). A common practice is to train a decoder based on neuronal activity patterns that were defined during awake periods, to identify similar ‘virtual trajectories’ during sleep^19,39,40^. In contrast to this approach, we used activity patterns defined during sleep to decipher the behavior of awake animals, which allows decoding performance to be validated against an actual observed behavior. Overall, these results demonstrate that the internal structure of neuronal activity reflects computational properties inherent to a given neural circuit.

Recent studies have shown that different visits to the same familiar environment are encoded by different subsets of hippocampal neurons^9,29^. Consequently, it has been hard to reconcile how memories can be stably stored over the long term in a circuit that yields an ever-changing neural code. Our findings that the internal structure of neuronal activity remains stable over time raises the possibility that stability of the neural code is achieved, in part, through a stable relationship between neuronal population activity patterns. Such a stable relationship may also support the encoding of new information via its integration into a pre-existing structured code. We also demonstrated the conservation of the internal structures of neuronal activity between mice. This conservation allowed us to infer the position or head direction of a mouse, based on the mapping between the behavior and activity patterns in another mouse. Thus, the analogy between the internal structures allows the meaning of network states to be exported from one animal to another (up to the symmetry of the internal structure).

The internal structure of neuronal activity highlights general features of the computational task executed by a brain circuit. Consistent with this idea, recent studies have shown that the same neural circuit may similarly encode different external variables in different behavioral tasks^41–43^. For instance, non-spatial representations in the entorhinal cortex form a hexagonal grid-like pattern^42,43^, similar to the known spatial tuning of grid cells^3,4,44^. Since analogous tuning properties were observed in both cases, it is predicted that they would have the same internal structure of neuronal activity—a torus^45,46^. Thus, the internal structure of neuronal activity serves as a fingerprint of the computations carried out by different neural circuits, enabling to investigate their function, and even redefine their identity.

## Acknowledgments

Y.Z. is a CIFAR-Azrieli Global Scholar in the Brain, Mind & Consciousness program, and an incumbent of the Daniel E. Koshland Sr. career development chair. Y.Z. is supported by grants from the Abraham and Sonia Rochlin Foundation, the Hymen T. Milgrom Trust, the Minerva Foundation, the Israel Science Foundation (grant 2184/14), the European Research Council (ERC-StG 638644), and the FP7 Marie Curie actions (CIG 630852). We thank Timothy O’Leary, Jerome Lecoq, Ofer Yizhar, Rony Paz, Nachum Ulanovsky, Yadin Dudai, Misha Tsodyks, Rafi Malach, Michal Schwartz, Michal Rivlin, Yoram Burak, and members of the Ziv lab for helpful advice and comments on the manuscript. We also thank Adrien Peyrache and Gyӧrgy Buzsáki for sharing their data sets.

## Supplementary materials

### Materials and methods

#### Animals and surgical procedures

All procedures were approved by the Weizmann Institute IACUC. We used male mice, aged 8-12 weeks at the beginning of the study. Mice designated for calcium imaging in the hippocampal CA1 were housed with 1-4 cage-mates, while mice used for imaging in the ACC were single housed. All cages contained running wheels. All surgical procedures were conducted under isoflurane anesthesia (1.5-2% volume). For hippocampal imaging, we used C57BL/6 wild type mice, which underwent two surgical procedures: virus injection and glass tube implantation. We injected 400 nL of the viral vector AAV2/5-CaMKIIa-GCaMP6f^47^ (~2 × 10^13^ particles per ml, packed by University of North Carolina Vector Core) to the CA1 at stereotactic coordinates: −1.9 mm anterio-posterior, −1.4 mm mediolateral, −1.6 mm dorsoventral relative to the Bregma. The injected mice were allowed to recover in their home-cages for at least 1 week before the subsequent surgical procedure. We next implanted a glass guide tube directly above the CA1, as previously described^9,29^. For ACC imaging, we used CaMKII-tTA and TRE-GCaMP6s double transgenic mice (Jackson stock no 003010 & 024742; referred to as CaMKII-GCaMP6), bred on a C57BL/6 background. They were implanted directly with a micro-prism lens (800μm diameter) in the ACC. Stereotactic coordinates of the implantation were: 1 mm anterior-posterior, 0 mm mediolateral (measured relative to the medial side of the prism), −1.8 mm dorsoventral from bregma.

#### Ca^2+^ imaging and behavioral setup

##### Preparatory process

For time-lapse imaging in freely behaving mice using an integrated miniature fluorescence microscope (nVistaHD, Inscopix), we followed a previously established protocol^9,29^. Briefly, at least 3 weeks after the surgical implantation procedure, we examined Ca^2+^ indicator expression and tissue health by imaging mice under isoflurane anesthesia using a two-photon microscope (Ultima IV, Bruker, Germany), equipped with a tunable Ti: Sapphire laser (Insight, Spectra Physics, Santa Clara, CA). For the CA1 implanted mice, we inserted into the guide tube a ‘microendoscope’ consisting of a single gradient refractive index lens (0.44 pitch length, 0.47 NA, GRINtech GmbH, Germany). We selected for further imaging only those mice that exhibited homogenous GCaMP6 expression and healthy appearance of the tissue. For the selected CA1 implanted mice, we affixed the microendoscope within the guide tube using ultraviolet-curing adhesive (Norland, NOA81, Edmund Optics, Barrington, NJ). Next, we attached the microscope’s base plate to the dental acrylic cap using light cured acrylic (Flow-It ALC, Pentron, Orange, CA). All mice were returned to their home cages for several days following the aforementioned procedure.

##### Calcium imaging in freely behaving mice

We trained the mice to run back and forth on an elevated 96cm linear track^9^. To record mouse behavior, we used an overhead camera (DFK 33G445, The Imaging Source, Germany), which was synchronized with the integrated microscope. Before initiating Ca^2+^ imaging, we trained the mice for 3–11 days. Training and imaging sessions consisted of five to seven 3-min-long trials, with an inter-trial interval of 3 minutes. Ca^2+^ imaging was performed at 20Hz or 10Hz in CA1 or ACC, respectively.

#### Processing of Ca^2+^ imaging data

We processed imaging data using commercial software (Mosaic, version 1.1.1b, Inscopix) and custom MATLAB routines as previously described^9,29^. To increase computation speed, we spatially down-sampled the data by a factor of two in each dimension (final pixel size of 2.3 × 2.3μm). To correct for non-uniform illumination both in space and time, we normalized the images by dividing each pixel by the corresponding value from a smoothed image. The smoothed image was obtained by applying a Gaussian filter with a radius of 100μm to the movies. Normalization also enhanced the appearance of the blood vessels, which were later used as stationary fiducial markers for image registration. We used a rigid-body registration to correct for lateral displacements of the brain. This procedure was performed on a high contrast subregion of the normalized movies at which the blood vessels were most prominent. The movies were transformed to relative changes in fluorescence, 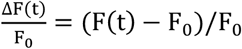, where F_0_ is the value for each pixel averaged over time. For cell detection, the movies were down-sampled in time by a factor of five. We detected spatial footprints corresponding to individual cells using an established cell detection algorithm that applies principal and independent component analyses (PCA and ICA). For each spatial footprint, we used a threshold of 50% of the footprint’s maximum intensity, and each pixel that did not cross the threshold was set to zero. After the cells were detected, further cell sorting was performed to identify the spatial footprints that follow a typical cellular structure. This was done by measuring the footprint area and circularity, and discarding those whose radius was smaller than 5μm or larger than 15μm, or which had a circularity smaller than 0.8. In some cases, the output of the PCA/ICA algorithm included more than one component that corresponded to a single cell. To eliminate such occurrences, we examined all pairs of cells with centroid distances < 18μm and whenever their traces had a correlation > 0.9, the cell with the lower average event peak amplitude was discarded. To identify the same neurons across multiple imaging sessions, we used a probabilistic method for cell registration^48^, which estimates the probability of correct registration for each cell in the data set, and the overall rates of registration errors.

#### Detection of Ca^2+^ events

Ca^2+^ activity was extracted by applying the thresholded spatial footprints to movies at full temporal resolution (20Hz) ΔF(t)/F_0_. Baseline fluctuations were removed by subtracting the median trace (20 sec sliding window). The Ca^2+^ traces were smoothed with a low-pass filter with a cutoff frequency of 2Hz. Ca^2+^ candidate events were detected whenever the amplitude crossed a threshold of 4 or 5 median absolute deviations (MAD), for GCaMP6s or GCaMP6f, respectively. We considered for further analysis only candidate Ca^2+^ events with indicator decay time for GCaMP6s or GCaMP6f equal to or longer than 600msec or 200msec, respectively. In order to avoid the detection of several peaks corresponding to a single Ca^2+^ event, only peaks that were 4 or 5 MAD higher than the previous peak (within the same candidate event) and 2 or 2.5 MAD higher than the next peak, for GCaMP6s or GCaMP6f, respectively, were regarded as true events. We set the Ca^2+^ event occurrence to the time of the peak fluorescence. To mitigate the effects of crosstalk (i.e., spillover of Ca^2+^ fluorescence from neighboring cells), we adopted a conservative approach, allowing only one cell from a group of neighbors (pairs of cells with centroid distances < 18μm) to register a Ca^2+^ event in any 200msec time window. If multiple Ca^2+^ events occurred within ~200msec in neighboring cells, we retained only the event with highest peak ΔF(t)/F_0_value. If two neighboring cells had a correlation > 0.9 in their events, the cell with the lower average peak amplitude was discarded. After the events were identified, further event sorting was performed to find the cells with sufficient signal-to-noise ratios. This was done by measuring the event rate and the average event peak amplitude for each cell and discarding those whose event rate was smaller than 0.01Hz or which had an average event amplitude smaller than 1% (ΔF(t)/F_0_). We considered each neuron to be active for two consecutive frames at the peak of each detected Ca^2+^ transient (to account for the typical Ca^2+^ indicator rise time).

#### Electrophysiology in the subiculum and thalamus

We obtained published electrophysiology recordings^20^ from multiple anterior thalamic nuclei, mainly the anterodorsal nucleus (ADn), and subicular areas, mainly the post-subiculum (PoS), in freely moving mice foraging for food in an open environment (53 × 46 cm). The authors recorded at 20kHz, simultaneously from 64 to 96 channels, and processed the raw data to extract the LFPs and detect spikes.

#### Obtaining the data points

For Ca^2+^ imaging data, we constructed a binary activity matrix of size NxK, where N is the number of neurons and K is the number of frames in the movies tracking Ca^2+^ dynamics. The i^th^ by j^th^ element of the matrix was set to 1 if the i^th^ neuron was active in the j^th^ frame, and was otherwise set to 0. We then defined each data point as a frame-level binary activity vector of length N, namely, a row of the activity matrix. For further analysis of the data, we used only frame-level activity vectors with >1 active neurons (non-zero elements).

For electrophysiology data, we first binned the activity of the neurons using a 100msec time bin. We then constructed a binary activity matrix of size NxK, where N is the number of neurons and K is the number of time bins. The i^th^ by j^th^ element of the matrix was set to 1 if the i^th^ neuron fired at least one time during the j^th^ time bin, and was otherwise set to 0. We then defined each data point as a bin-level binary activity vector of length N, namely, a row of the activity matrix. For dimensionality reduction and further analysis of the data, we discarded activity vectors with <15 active neurons.

#### Dimensionality reduction

Non-linear dimensionality reduction techniques enable identification of sets of activity patterns which lay in a low dimension manifold, within the high dimension of the data (N, number of neurons), even if this manifold is non-linear (**Supplementary Fig. 1**). To this end, we used Laplacian Eigenmaps (LEM) for non-linear dimensionality reduction as previously described^11^. In general, such techniques utilize the local relationships between proximal data points to reconstruct a global distance metric. Given K data points x1,…, xK (number of frames) lying in an N-dimensional space of neuronal activity, we constructed a weighted graph with K nodes, one for each point, and a set of edges connecting neighboring data points. We considered nodes i and j to be connected by an edge if i is among p of the nearest neighbors of j, or j is among p of the nearest neighbors of i (p=0.25-0.5% for all data sets). We used the parameter free (“simple minded”) method for choosing the weights, i.e., W_ij_ =1 if nodes i and j are connected, and 0 otherwise ^11^. We then computed the eigenvalues and eigenvectors for the generalized eigenvector problem: Lf = λDf, where D is the diagonal weight matrix, and its entries are column sums of W, and L = D − W is the Laplacian matrix. We left out the leading eigenvector (as previously described^11^) and used the next 10 eigenvectors for embedding in a 10-dimensional Euclidean space. The LEM was performed twice. For the second iteration, performed on the reduced dimensional data, we defined nodes i and j as connected if i is among p of the nearest neighbors of j, or j is among p of the nearest neighbors of i (p=7.5-15% for all data sets). For further analysis, we used the three leading eigenvectors (after leaving out the first eigenvector).

#### Estimation of data internal dimensionality and topology

To estimate the internal dimension of the data, for any given data point in the reduced space, we calculated the number of neighbors within a sphere surrounding it, as a function of the sphere’s radius. We used the slope of the number of neighboring data points within a given radius on a log-log scale to estimate the dimension of the data^14^. A simulation illustrating this procedure is presented in **Supplementary Fig. 2**.

To estimate the topology of the data, we calculated the numbers of components (β_0_), holes (β_1_), and spaces (β_2_), as a function of the radius threshold, using a previously established algorithm^23^ (Javaplex: A research software package for persistent (co) homology, https://github.com/appliedtopology/javaplex). We then searched for components, holes, and spaces that were stable across a wide range of radii. A simulation illustrating this procedure is presented in **Supplementary Fig. 7**. Since the algorithm is computationally demanding and is not scalable to large data sets, we applied the algorithm to a representative set of data points. To obtain this representative set, we applied a clustering procedure (K-means) on the reduced dimensional data and extracted the centroids of the clusters. To overcome sensitivity to noise, we discarded sparse clusters (<50 data points). To ensure a final number of cluster centroids >50, we used K=70 for the K-means procedure.

#### K-means clustering

In cases in which the data points are continuously distributed, there is no natural separation of the data points into discrete clusters. Therefore, for parameterization of the encoded variable, we used K-means, which yielded segmentation of the data into a discrete set of compact subsets. For electrophysiology data, we used K-means (K=8) to segment the data into different clusters.

#### Topological clustering and sub-clustering

In cases in which the distribution of data points consists of a discrete set of dense clusters, these clusters can be considered as components in terms of data-topology, and the same algorithm that estimates the number of components (β_0_, see estimation of data internal topology) naturally yields a topological-based clustering of the data points. To utilize the topological approach for data clustering, we focused on large components (>250 data points for clustering and >50 data points for sub-clustering), and chose a radius that captures the maximal number of stable components (see estimation of data internal topology). To assign data points that do not belong to one of the defined components, we associated each unassigned data point with the cluster that contains its nearest assigned neighbor. This procedure was performed by gradually increasing the radius threshold, while not changing the assignment of previously assigned data points.

#### Temporal segmentation of the data

For the Ca^2+^ imaging data from the CA1 and ACC, we used the temporal sequence of clusters to temporally segment the data. Applying a segmentation procedure to the data points allowed us to define the beginning and end of each time segment, and at the same time, reduce the number of short deviations from a given cluster that could result from noise and sparse neuronal activity. To segment the data points, we defined two thresholds: (1) maximal interval between consecutive data points that belong to the same cluster; (2) minimal number of data points within a segment. To optimize the segmentation procedure, we systematically tested different values of the maximal interval threshold and examined the obtained distribution of segment lengths. A suitable threshold should result in a robust distribution (not sensitive to small changes of threshold value) of segment lengths. The distribution of number of data points in a candidate segment was clearly bimodal, which allowed valid segments to be distinguished from noise. We performed segmentation independently for each cluster enabling data points to belong to a single cluster, multiple clusters, or none of the clusters. In practice, the majority of data points belonged to a segment of only one cluster. After segmenting the data, we constructed an average activity vector by measuring the average activity of each neuron during a given time segment. For each set of segments from the same cluster, we applied principal component analysis (PCA) to the obtained average activity vectors to expose intra-cluster heterogeneity, corresponding to different segment subtypes within a given cluster.

#### Calculating the transition matrix

To capture the temporal relationship between the different network states, we calculated the transition matrix, i.e., the probability of a data point to belong to cluster i at time t+1 given that the preceding point belongs to cluster j at time t. Similar analysis was performed at the segment level where the transition matrix was defined by the probability of a segment to belong to cluster i given that the preceding segment belongs to cluster j.

#### Temporal ordering of clusters and sub-clusters

For the Ca^2+^ imaging data from the CA1 and ACC, we examined the internal structure of the data within a given cluster or segment subtype. We sub-clustered the reduced dimensional data points using topological clustering as described above. To evaluate the temporal ordering of the sub-clusters, for each possible ordering (i.e. permutation), we calculated the sum of probabilities of moving from each sub-cluster to its consecutive sub-cluster. The ordering 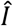 of the M sub-clusters was set as the order that maximized the sum of probabilities out of the M! possible permutations.

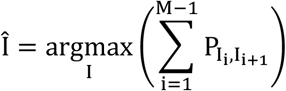

Since examining all possible permutations is computationally demanding, we first calculated the number of times that each sub-cluster appeared at the beginning and at the end of a segment. The analysis revealed that one sub-cluster was prominent at the start of a segment and another sub-cluster was prominent at the end of the segment, implying a specific directionality for the ordering of the sub-clusters. This allowed us to constrain the first and last sub-clusters among the M sub-clusters, and consequently we had to examine only (M-2)! possible permutations rather than M!.

For electrophysiology data from the ADn and PoS, due to the observed ring topology of the reduced dimensional data (**Supplementary Fig. 7**) and the trajectory within it (**Supplementary Fig. 8a-b**), we sought to cyclically order the obtained M clusters. To this end, for each ordering, we calculated the sum of probabilities for moving from a cluster to its neighboring clusters from both directions. The cyclical ordering of sub-clusters was set to maximize the sum of these probabilities, out of the (M-1)!/2 possible orders (due to rotation and reflection symmetry). The ordered states of neuronal activity allowed us to reconstruct the encoded variables and calculate the internal tuning curve for each cell.

#### Reconstruction of the encoded variables

To reconstruct the variables encoded within the network activity, we parameterized the network states. For Ca^2+^ imaging data from the CA1, we focused on the reconstruction of an internal position within clusters associated with locomotion. We used the obtained sub-clusters and their ordering, and assigned evenly spaced internal positions to the different sub-clusters. For comparison with the external position, we set the reflection degree of freedom (out of the two possible orientations of the linear track due to its symmetry) by minimizing the mean squared error between the internal reconstruction and the external variable (performed globally for the entire data set). The error (mismatch) between the estimated position (or phase) based on the reconstructed internal representation 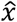, and the actual position (or phase)*x*, was defined as 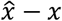.

For electrophysiology data from the ADn and PoS, we reconstructed the internal angle based on the clustering of the entire data set, and the obtained cyclical ordering of the clusters. Data points that belong to cluster k were assigned an internal angle of 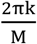, where M is the number of clusters. We then used a truncated Gaussian kernel W (σ = 2 frames, size = 5 frames) to temporally smooth the data. For comparison with the head direction, we set the rotation and reflection degrees of freedom (due to the symmetries of a ring) by minimizing the mean squared error between the internal reconstruction and the external variable (performed globally for the entire data set). The mismatch between the estimated head direction based on the reconstructed internal representation 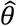, and the actual head direction θ, (or the decoded head direction during sleep; see *REM sleep decoder*) was defined as 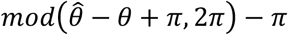.

#### Calculation of internal tuning curves

We sought to calculate the tuning curve for each cell relative to the state of the network, rather than relative to any external variable (e.g., position). For this calculation, we first measured the time the network spent (occupancy) at any given state and the number of neuronal activity events within each network state (Ca^2+^ event number or number of spikes). We then divided the number of neuronal activity events by the occupancy, obtaining the activity rate as a function of the internal state of the network – hence the internal tuning curve.

For Ca^2+^ imaging data from the CA1 and ACC, we focused on the reconstruction of internal tuning curves within clusters associated with locomotion. Internal tuning curves were calculated for cells that displayed >5 Ca^2+^ events within the relevant clusters. For comparison with external tuning curves, after we calculated the occupancy and Ca^2+^ event number vectors, we interpolated the Ca^2+^ event number and the occupancy at the ordered network states (sub-clusters) to increase the number of states so it will match the number of spatial bins. We then used a truncated Gaussian kernel (σ = 1.5 bins, size = 5 bins, as used for calculation of external tuning curves) to smooth the two functions. Finally, we computed the internal tuning curve for each neuron by dividing the smoothed map of Ca^2+^ event numbers by the smoothed map of occupancy.

For electrophysiology data from ADn and PoS, the internal tuning curves were calculated based on the clustering of the entire data set, and the obtained cyclical ordering of the clusters. Data points were smoothed as described above (see *Reconstruction of the encoded variables*). We then binned the internal angle into 40 bins of 9° and calculated the occupancy at any given angular bin and the neuron’s number of spikes within each bin. Finally, we computed the internal tuning curve for each neuron by dividing the number of spikes by the occupancy.

#### Calculation of external tuning curves

For Ca^2+^ imaging data from the CA1 and ACC, we calculated the tuning of cells to location. We considered periods wherein the mouse ran > 1cm/sec. We divided the track into 24 bins (4cm each), and excluded the last 2 bins at both ends of the tracks where the mouse was generally stationary. We computed the occupancy and the number of Ca^2+^ events in each bin, and then smoothed these two maps (occupancy and Ca^2+^ event number) using a truncated Gaussian kernel (σ = 1.5 bins, size = 5 bins)^9,29^. We then computed the activity map (event rate per bin) for each neuron by dividing the smoothed map of Ca^2+^ event numbers by the smoothed map of occupancy. For hippocampal data we separately considered place fields for each of the two running directions on the linear track. For data recorded from the ACC, we pooled the data from both running directions on the linear track while taking into account the running phase rather than the absolute location. The pooling was performed by flipping the positional indexing of the linear track while the mouse was running in a given direction (this way, a given position on the track while the mouse was running to the right corresponded to the mirror location while the mouse was running to the left).

For the electrophysiology data from the ADn and PoS, we computed the tuning to head direction. We first binned head directions into 40 bins of 9°. We then calculated for each neuron the average firing rate given an angular bin, by dividing the number of spikes in each bin by the occupancy within that bin.

#### Comparison between internal and external tuning curves

For Ca^2+^ imaging data from the CA1 and ACC, we measured the mismatch between the internal and external tuning curves. For CA1 data the mismatch was defined as the difference between the internal and external preferred position, and for the ACC it was defined as the difference between the internal and external phase. To test the significance of the similarity in the coding properties, we compared the average absolute mismatch to that obtained for 1,000 shuffled data sets. For each of the 1,000 shuffled sets, we shuffled between the identities of the cells.

For the electrophysiology data from the ADn and PoS, we quantified the angular tuning of each neuron and compared between the internal and external tuning curves. To quantify the angular tuning, we calculated for each angular tuning curve (either to internal angle or to head direction) the Rayleigh vector: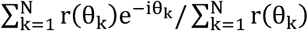, where N is the number of evenly spaced angular bins, 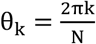, and r(θ) is the firing rate of the neuron given the angle θ. The preferred direction of a neuron was estimated by the angle of the Rayleigh vector of its tuning curve, and the directionality (the degree of its tuning to a single direction) was estimated by the absolute value (length) of the Rayleigh vector. For both preferred direction and directionality (Rayleigh vector length) we compared the values obtained for each neuron from the internal and external tuning curves. To further compare the internal and external tuning curves of each neuron, we calculated the Pearson correlation between the two tuning curves. For the visualization of the data in **Supplementary Fig. 8e-g**, we labeled cells as head-directions cells if they had Rayleigh vector length of >0.5 and peak firing rate >5Hz.

In both cases, in order to compare the internal tuning curves to the external tuning curves, we used the same degrees of freedom obtained in the *Reconstruction of the encoded variables* (described above).

#### REM sleep decoder

For the electrophysiology data from the ADn and PoS, we calculated the ‘virtual trajectory’ of the internal angle during REM sleep periods as described above (see *Reconstruction of the encoded variables*). Since the actual head direction is constant during sleep, the internal angle was compared to the ‘virtual trajectory’ of head direction obtained by a maximum likelihood decoder. The maximum likelihood decoder for virtual head direction during REM sleep was based on the external tuning curves during wake periods, using only head direction cells (>0.5 Rayleigh vector length and >5Hz peak firing rate). For comparison between the internal angle and the decoded head direction, we sought the rotation and reflection degrees of freedom (due to the symmetries of a ring) that minimize the decoding mean squared error.

#### REM sleep to wake decoder

We trained a decoder during periods of REM sleep and tested it during periods of awake behavior. The internal angle and internal tuning curves were first calculated for REM sleep data, as described above (see *Calculation of internal tuning curves*), and were then used to train a maximum likelihood decoder (**Fig. 4f**). Next, the decoder was tested on neuronal activity form awake periods. For comparison of the decoded head direction with the actual head direction, we sought the rotation and reflection degrees of freedom (due to the symmetries of a ring) that minimize the decoding mean squared error (inset of **Fig. 4f**). The decoding error was compared to that obtained for shuffled data (see *Shuffle test*).

#### Across-mice decoder

Across-mice decoders rely on the analogy between the internal structures of neuronal activity across different mice (or different days of the experiment) for the same brain region. These decoders were used to infer the external state (position or head direction) of one mouse based on the relations between internal state (neuronal activity patterns) and external state of another mouse (or of the same mouse at another day of the experiment). We used the neuronal activity within the reduced dimensional space to parametrize data points within the internal state. Specifically, for the Ca^2+^ imaging data, clusters and ordered sub-clusters therein were translated into position on a 1D segment. For electrophysiology data, we used the circular order of the clusters to assign an angle to each data point. Based on this parametrization, we defined mapping across network states within internal structures obtained for different mice (or different days for the same mouse). In all cases, the mapping was defined up to the internal symmetry of the internal structure: rotation and reflection for PoS and ADn data and reflection for the CA1 data. For mouse 1, neuronal activity at each time frame was associated with a specific network state. Based on the similarity between internal structures, an analogous network state is found in mouse 2. This network state of mouse 2 is associated with a known external state. The decoded external state of mouse 1 was set as the same associated external state of mouse 2. The external state decoding error was calculated and compared to that obtained for shuffled data (see *Shuffle test*).

#### Phase decoder

For Ca^2+^ imaging data recorded from the ACC, we tested the similarity between neuronal activity during running in one direction of the linear track and the neuronal activity during running in the other direction. We first defined the trajectory phase as the distance of the mouse from the starting point, namely, the distance from the left edge when running rightward and the distance from the right edge when running leftward. We then constructed a decoder that estimates the trajectory phase in a given direction (test data) based on the relations between neuronal activity pattern and trajectory phase in the other direction (training data). Specifically, for each data point obtained during running in a given direction, we found its nearest neighbor in the reduced space of neuronal activity among all data points obtained during running in the opposite direction. The decoded phase was set as the animal’s trajectory phase at the time of the nearest neighbor data point. The trajectory phase decoding error was calculated and compared to that obtained for shuffled data (see *Shuffle test*).

#### Shuffle test

To test the significance of decoding accuracy, we measured the difference between the decoded values and the measured behavior at each time bin. We then compared the performance of each of the decoders described above to the performance obtained for 1,000 shuffled data sets. For each of the 1,000 shuffled sets, we shuffled the decoded values across the different time bins, and set the degrees of freedom (rotation and reflection for angle, and reflection for position) to those that minimized the decoding mean squared error (done separately for each shuffle).

#### Labeling of behavioral states

For Ca^2+^ imaging data from the CA1 and ACC, we manually labeled the behavior of the mouse. To identify periods corresponding to behavioral states of “Running”, “Drinking”, “Turning”, and “Rearing”, we analyzed the movies from the overhead camera that tracked animal behavior using video tracking software (EthoVision XT 11.5). We manually tagged the beginnings and ends of each behavioral state throughout each movie. Further separation of the running state to the start and end of running was done based on animal position along the linear track: “Start run” and “End run” were defined from the starting point to the center of the linear track, and from the center to the end point of the track, respectively. This behavioral labeling was used both for the comparison with neuronal clustering (**Fig. 2f**), and for the calculation of correlations between representations of behavioral states (**Supplementary Fig. 6**).

#### Statistical analysis

Statistical analysis was conducted using MATLAB 2016b software. One-tailed unpaired t-test with Holm–Bonferroni correction for multiple comparisons was conducted for between-regions comparisons of activity correlation level.

**Supplementary Fig. 1:**
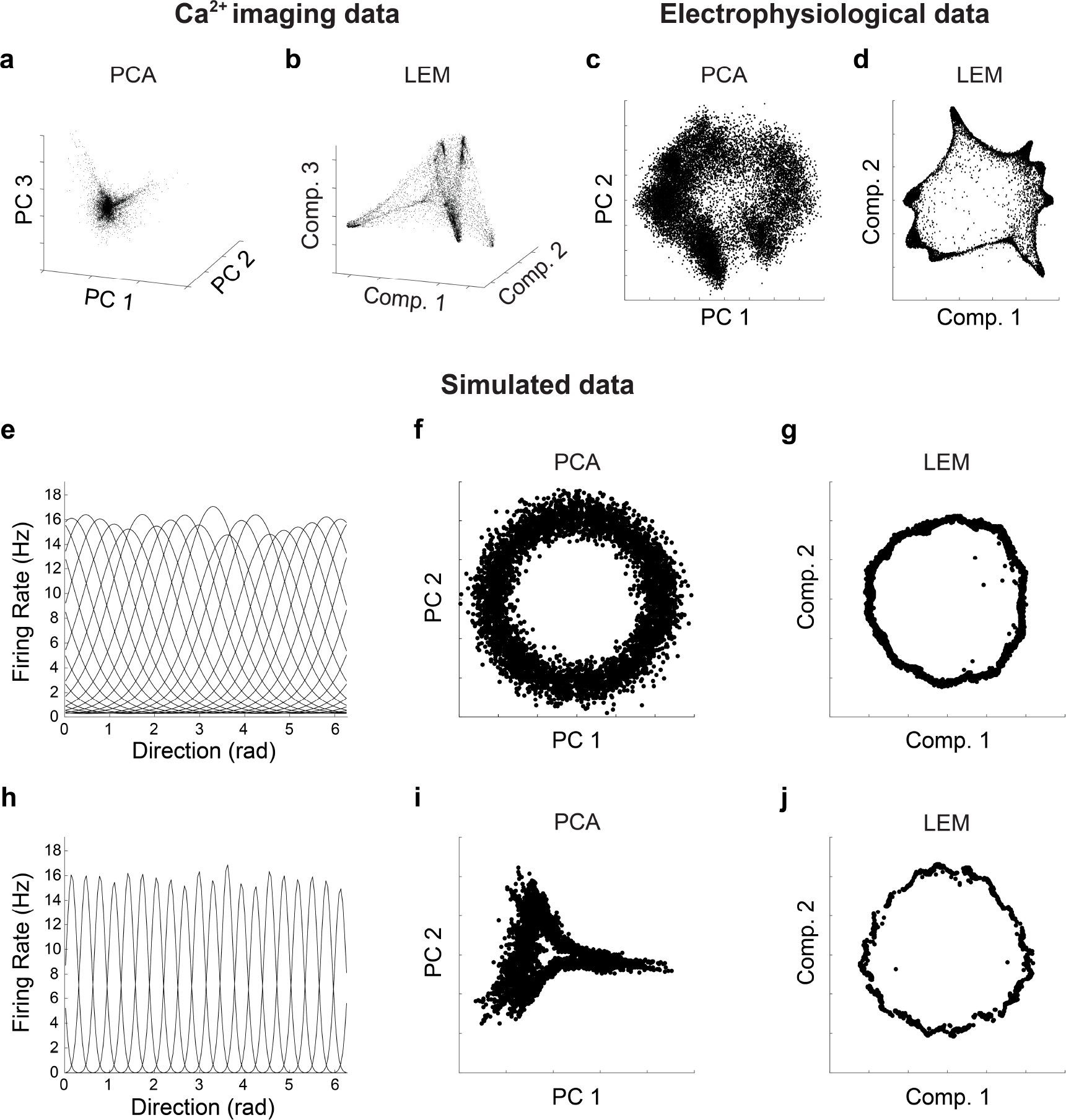
Non-linear dimensionality reduction enables more accurate estimation of internal structure than linear methods. (**a-b**) Ca^2+^ imaging data: the distribution of data points in the reduced dimensional space of neuronal activity, using PCA and (**a**) and LEM (**b**). (**c-d**) Electrophysiological data: the distribution of data points in the reduced dimensional space of neuronal activity, using PCA and (**c**) and LEM (**d**). (**e** and **h**) We simulated neuronal responses for 20 conditionally independent neurons which follow Poisson statistics. The centers of the tuning curves were equally spaced on a one-dimensional circular variable, and were either widely tuned (**e**) or narrowly tuned (**h**). (**f-g**) The distribution of data points in the reduced space of neuronal activity, using PCA (**f**), and LEM (**g**), for widely tuned neurons. Both methods captured the ring topology of the data. (**i-j**) The distribution of data points in the reduced space of neuronal activity, using PCA (**i**), and LEM (**j**), for narrowly tuned neurons. LEM exposed the ring topology more successfully compared to PCA.

**Supplementary Fig. 2:**
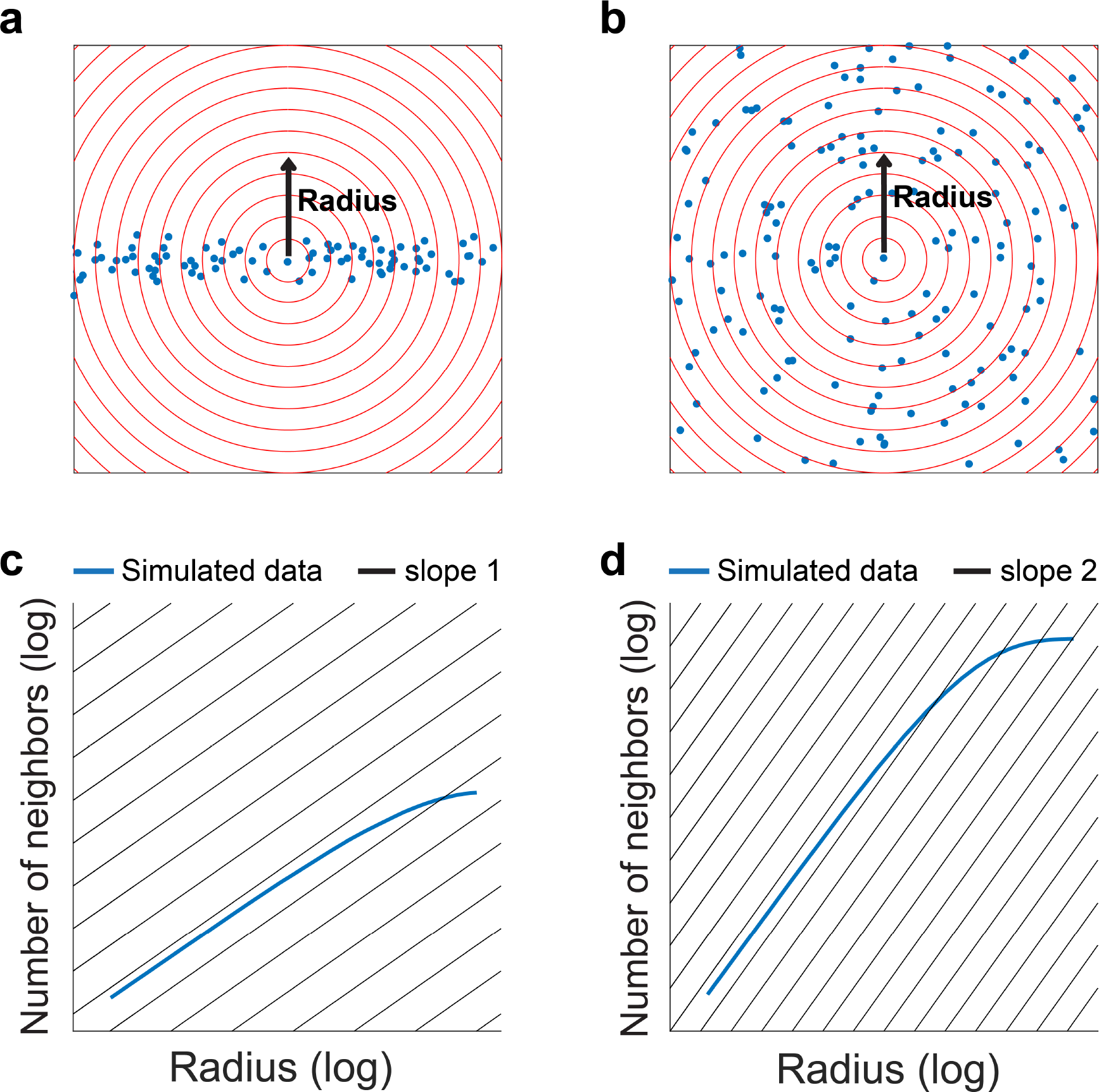
Estimation of the internal dimension of the data. (**a-b**) Simulated data points, for 1-dimensional data (**a**) and for 2-dimensional data (**b**). The red circles indicate different distances (radii) from a given data point. (**c-d**) The average number of neighboring data points increases with the radius obeying a power law. The power is indicative of the internal dimension of the data. The number of neighboring data points increases linearly (**c**) or quadratically (**d**) with the radius, for 1-dimensional and 2-dimensional data, respectively.

**Supplementary Fig. 3:**
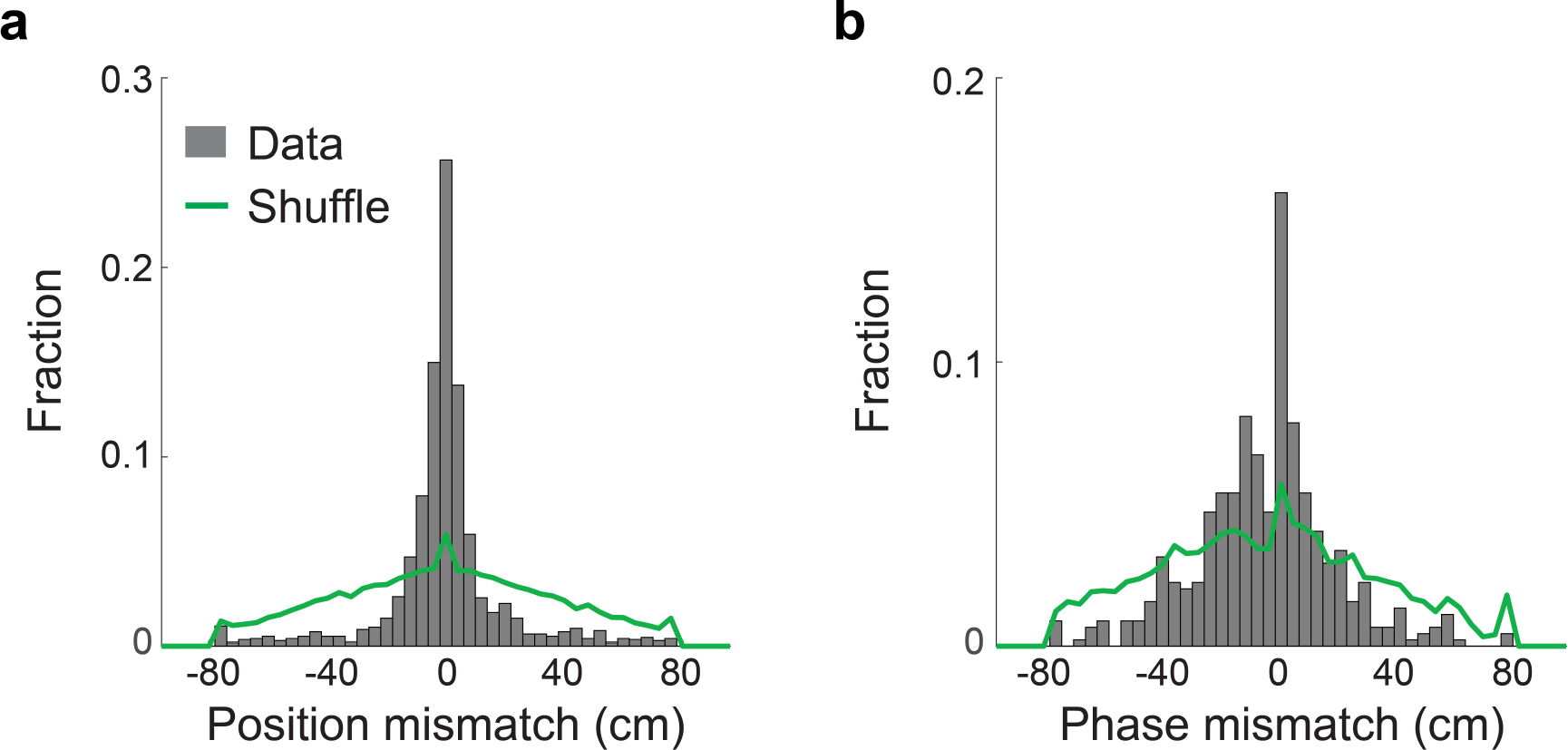
Internal tuning curves of the hippocampal CA1 and ACC reveal similar coding properties to those of their corresponding external tuning curves. (**a-b**) The distribution of differences in the preferred position (**a**) and preferred phase (**b**) between the calculated internal and external tuning curves of the same neurons, for data recorded from the hippocampal CA1 (**a**) and from the ACC (**b**). Data are represented in gray, and shuffled data in green. Data in **a** pooled from N=4 mice, and in **b** from N=3 mice.

**Supplementary Fig. 4:**
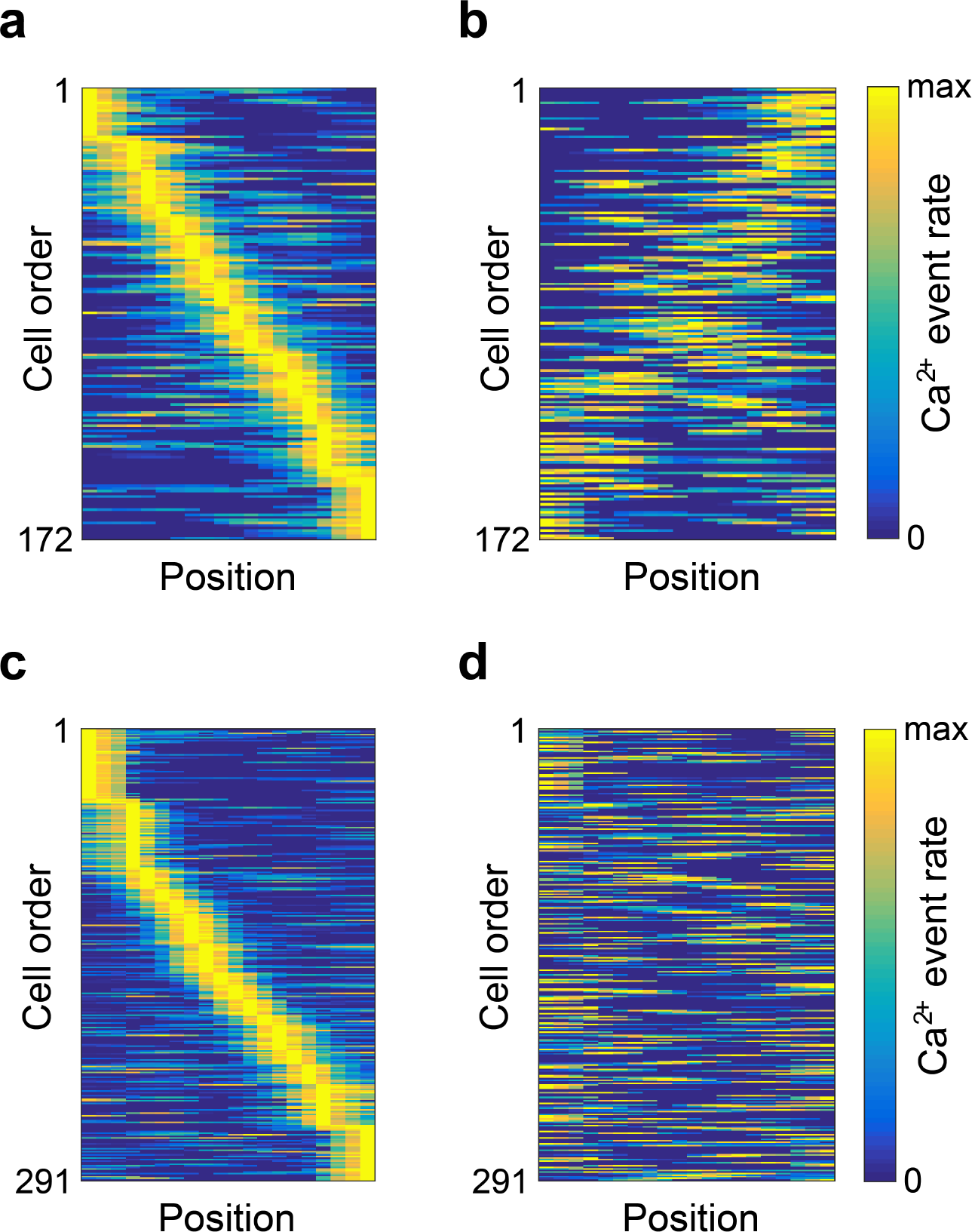
Neurons in the ACC, but not in the CA1, encode the trajectory phase for mice running on a linear track. For each neuron, we calculated its event rate as a function of the animal's position (spatial tuning curves) when running in a given direction. (**a**) Spatial tuning curves for ACC neurons ordered according to their preferred position along the track. (**b**) Spatial tuning curves for the same neurons with the same order as in **a**, calculated for the opposite running direction. (**c**) Spatial tuning curves for hippocampal neurons, ordered according to their preferred position along the track. (**d**) Spatial tuning curves for the same neurons with the same order as in **c**, calculated for the opposite running direction. Note that while in the hippocampus there is no clear relationship between the preferred positions of the same neurons in the two different running directions, the neurons in the ACC tend to have a symmetrical preferred position for the two directions, hence coding the trajectory phase.

**Supplementary Fig. 5:**
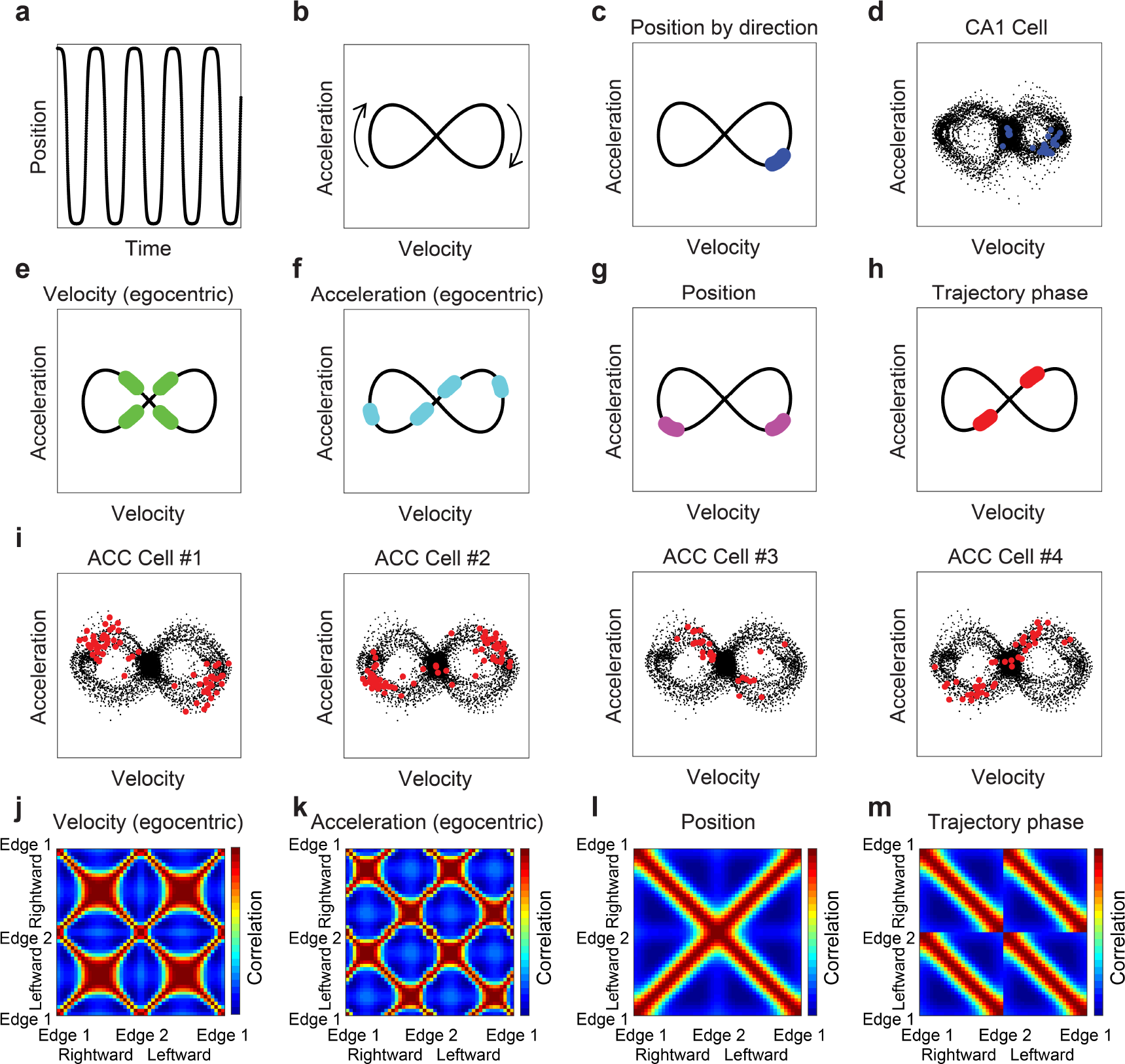
The encoding of trajectory phase in the ACC could not be accounted for by velocity or acceleration. (**a**) A simulated trajectory along a linear track. (**b**) The acceleration versus the velocity for the trajectory in **a**. (**c**) The activity of a simulated place cell (blue dots) which is responsive at a particular location in one running direction, overlaid on the velocity-acceleration plot in **b**. (**d**) The activity of an example place cell from the hippocampal data (blue dots), overlaid on the velocity-acceleration of the mouse (black dots). (**e-h**) The activity of a simulated cell overlaid on velocity-acceleration plot, for a cell that is responsive to a given speed (**e**), acceleration (**f**), position (**g**), or trajectory phase (**h**). (**i**) The activity of four example cells from the ACC data (red dots), overlaid on the velocity-acceleration of the mouse (black dots). The activity matches the simulated trajectory phase cells. (**j-m**) Pearson correlation between simulated ensemble activity patterns from different spatial locations on the linear track, for cells that are encoding speed (**j**), acceleration (**k**), position (**l**), or trajectory phase (**m**). The ensemble activity (as shown in **Fig. 2m**) matches the simulated trajectory phase ensemble.

**Supplementary Fig. 6:**
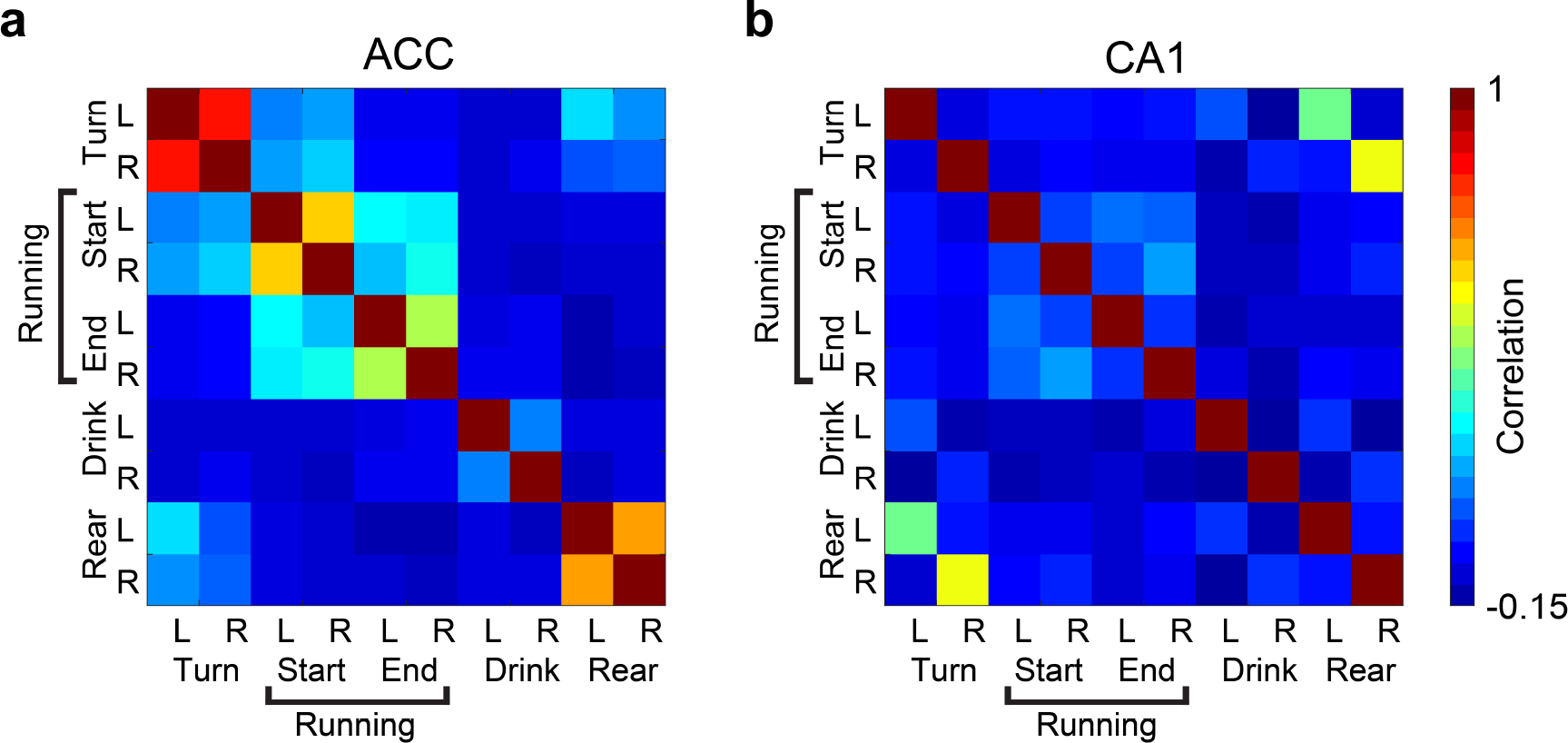
Neuronal activity during epochs of a given behavioral state are similar across opposite sides of the track, for data recorded from the ACC, but not from the hippocampus. (**a-b**) Pearson correlation between ensemble activity patterns given the behavioral state and side of the linear track, for the ACC (**a**) and hippocampal CA1 (**b**). Ensemble activity patterns are defined as the mean event rate for each neuron given a behavioral state on the same side of the track or the same running direction. L – left side or leftward epochs, R – right side or rightward epochs. Data averaged over N=3 mice in the ACC, and N=4 mice in the hippocampus.

**Supplementary Fig. 7:**
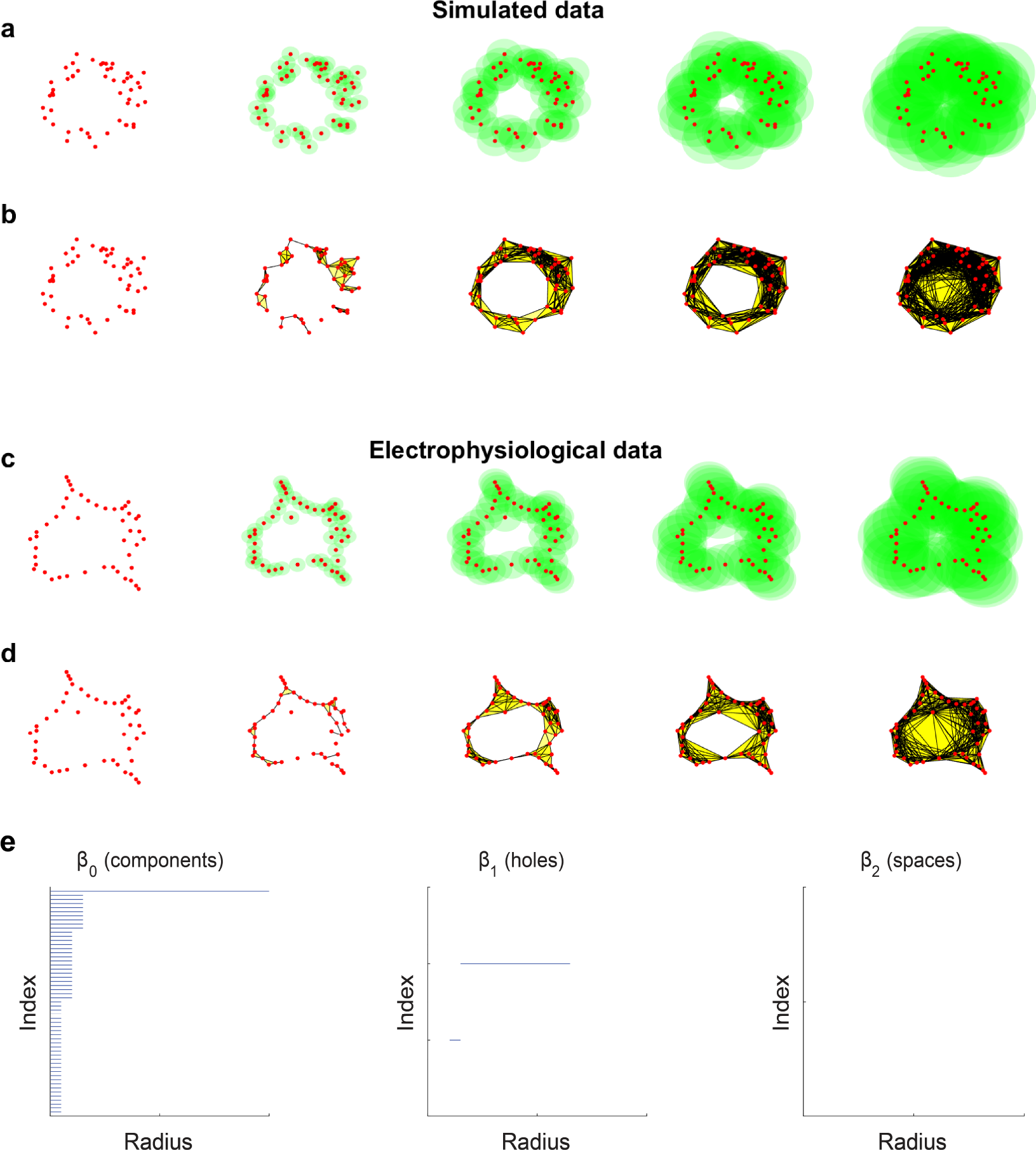
Measuring the internal topology of the data based on the estimation of the numbers of components, holes, and spaces. (**a-b**) Simulated data with a ring topology (**a**) Simulated data points (red dots) and an increasing radius (from left to right) represented by filled green circles around each data point. (**b**) The connections (black lines) between data points with overlapping surrounding spheres for different radii of the sphere (increasing from left to right). Triplets (cliques of three data points that are all interconnected) are marked by yellow areas. (**c-d**) Head direction data. (**c**) Representative data points (centers of mass of clustered data, red dots) and an increasing radius (from left to right) represented by filled green circles around each data point. (**d**) The connections (black lines) between data points with overlapping surrounding spheres for different radii of the sphere (increasing from left to right). Triplets (cliques of three data points that are all interconnected) are marked by yellow areas. (**e**) Quantifying the internal topology of the electrophysiological data by estimating the number of separate components (β_0_), number of holes (β_1_), and number of spaces (β_2_). We observed one component, one hole, and zero spaces for a wide range of radii, as expected for a ring topology.

**Supplementary Fig. 8:**
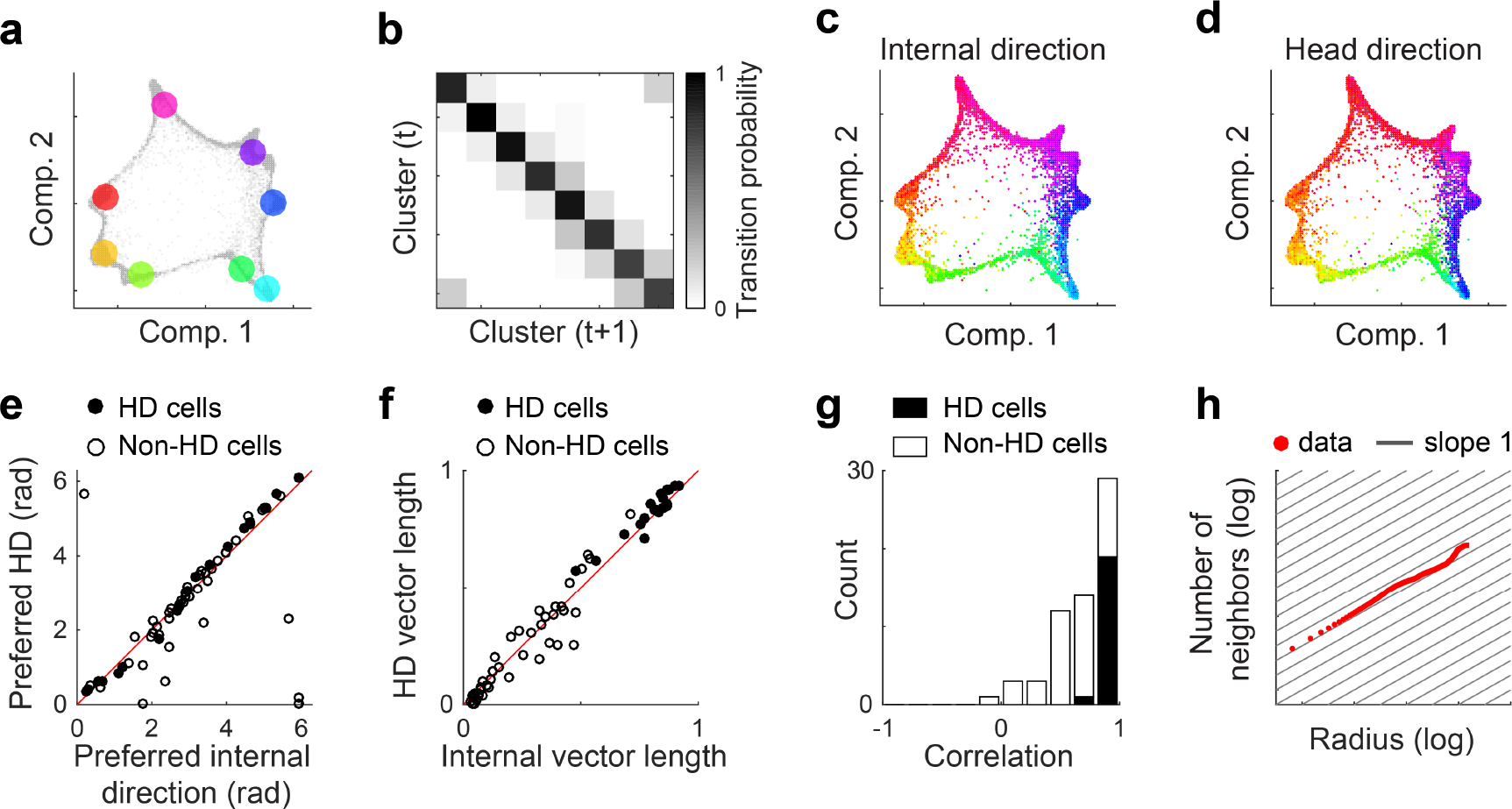
Internal representation of head direction obtained without relying on behavioral data. (**a**) Clustering of the data in **Fig. 4b**. The centers of masses of each cluster (colored dots) are overlaid on the data points in **Fig. 4b**(gray shaded dots). (**b**) The transition matrix (i.e., the probability of a data point appearing in cluster i, given that the preceding data point was in cluster j) shows that consecutive data points are more likely to be in the same or adjacent clusters. (**c-d**) The distribution of data points in the reduced dimensional space of neuronal activity, color coded according to the reconstructed internal direction (**c**), and the actual head direction (**d**). (**e-f**) The preferred direction (**e**), and Rayleigh vector length (**f**) calculated for the internal tuning curves (x-axis) and for the external tuning curves (y-axis). Filled circles indicate cells significantly tuned to head direction. (**g**) Distribution of correlations between internal and external tuning curves. Cells significantly tuned to head direction are shown in black. (**h**) Estimation of internal dimension: Cumulative number of neighboring data points as a function of the radius in the reduced dimensionality space, plotted on a log-log scale. The slope of the data (red) is approximately one (black lines), indicating one-dimensional data.

**Supplementary Fig. 9:**
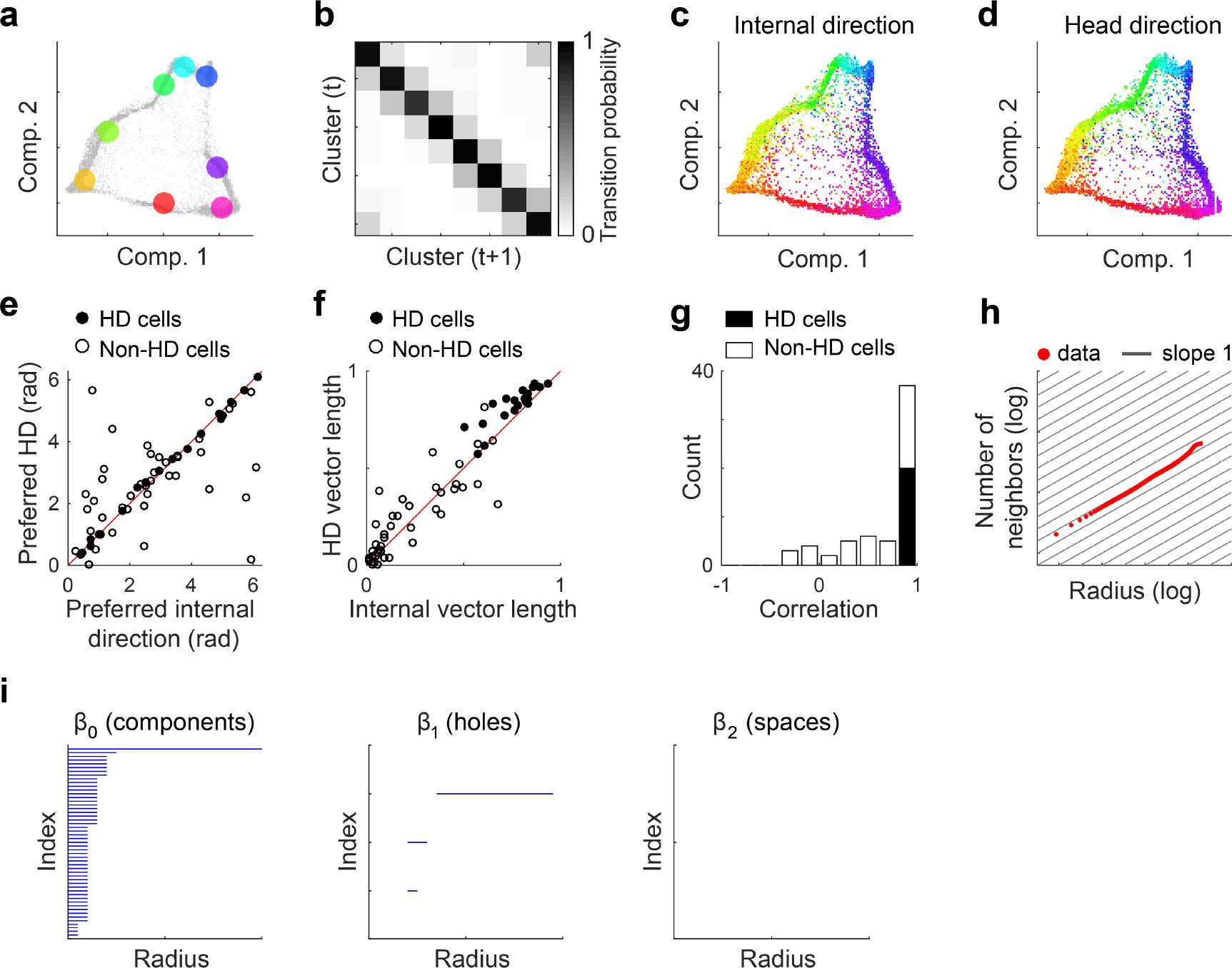
Internal representation of head direction during REM sleep. (**a**) Clustering of the data in **Fig. 4C**. The centers of mass of the clusters (colored dots) are overlaid on the data points in **Fig. 4C**(shaded gray). (**b**) The transition matrix (i.e., the probability of a data point to be in cluster i given that the preceding data points was in cluster j) shows that consecutive data points are more likely to be in the same or adjacent clusters. (**c-d**) The distribution of data points in the reduced dimensional space of neuronal activity color coded according to the reconstructed internal direction (**c**) and to the estimated head direction (**d**). (**e-f**) The preferred direction (**e**) and Rayleigh vector length (**f**) calculated for the internal tuning curves during REM sleep (x-axis) and for the external tuning curves during wake periods (y-axis). Filled circles indicate significantly tuned head direction cells. (**g**) Distribution of correlation between internal (during REM) and external (during wake) tuning curves. Significantly tuned head direction cells are shown in black. (**h**) Estimation of internal dimension: Cumulative number of neighboring data points as a function of the radius in the reduced dimensionality space, plotted on a log-log scale. The slope of the data (red) is approximately one (black), indicating one-dimensional data. (**i**) Measuring the internal topology of the data based on the estimation of the numbers of components (β_0_), holes (β_1_), and spaces (β_2_). Even during REM sleep we obtain one component, one hole, and zero spaces, for a wide range of radii. Overall these results indicate that the relationships between the neuronal activity patterns are preserved even in the absence of sensory input, reflecting inherent computational properties of these circuits.

### Supplementary Movie Legends

**Supplementary Movie 1: Neuronal activity in the hippocampal CA1 reveals a high density of data points within a small number of clusters.** The distribution of data points in the reduced dimensional space of neuronal activity from different view angles is shown.

**Supplementary Movie 2: Different states of neuronal activity in the hippocampal CA1 correspond to different behavioral states depending on their side of the linear track.** Concatenated epochs of mouse behavior, which correspond to a given neuronal activity state are shown, and the neuronal activity state and index of the epoch are specified.

**Supplementary Movie 3: Different states of neuronal activity in the ACC correspond to different behavioral states regardless of the side of the linear track.** Concatenated epochs of mouse behavior correspond to a given neuronal activity state. The neuronal activity state and index of the epoch are specified.

**Supplementary Movie 4: Neuronal activity in the ADn and PoS forms a ring topology in a reduced dimensional space, with a non-periodic continuous trajectory.** Top left: the trajectory of neuronal activity (moving dot) overlaid on the distribution of data points in the reduced dimensional space of neuronal activity. Bottom right: the actual head direction of the mouse, represented by the angle of the clock hand. Data points colored according to head direction.

**Supplementary Movie 5: Neuronal activity in the ADn and PoS forms a ring topology in a reduced dimensional space during periods of REM sleep.** The trajectory of neuronal activity (moving dot) overlaid on the distribution of data points in the reduced dimensional space of neuronal activity. The internal structure and the trajectory within it are maintained during REM sleep, when the head direction is mostly constant. Color represents the head direction as reconstructed by a maximum likelihood decoder that was trained on data from wake periods.

**Supplementary Movie 6: Across-mice decoder infers the behavioral state and position of the animal, based on the mapping between the behavior and activity patterns in the hippocampus of another animal.** Top left: The trajectory of neuronal activity (moving dot) for mouse 1 overlaid on the distribution of data points in the dimensionality reduced space of neuronal activity in mouse 1 (first two components). Top left: the corresponding trajectory of neuronal activity (moving dot) for mouse 1 overlaid on the distribution of data points in the dimensionality reduced space of neuronal activity in mouse 2 (first two components). Bottom left: Behavioral data from mouse 1. Bottom right: Reconstructed behavioral data of mouse 1 based on the mapping between the behavior and activity patterns.

